# CD169^+^ macrophages orchestrate plasmacytoid dendritic cell arrest and retention for optimal priming in the bone marrow of malaria-infected mice

**DOI:** 10.1101/2022.04.03.486895

**Authors:** Jamie Fried, Mahinder Paul, Zhixin Jing, David Fooksman, Grégoire Lauvau

**Affiliations:** Albert Einstein College of Medicine, Department of Microbiology and Immunology, Bronx, NY, USA, 10461; Albert Einstein College of Medicine, Department of Pathology

## Abstract

Plasmacytoid dendritic cells (pDC) are the most potent producer of type I interferon (IFN), but how pDC are primed *in vivo* is poorly defined. Using a mouse model of severe malaria, we have previously established that upon priming by CD169^+^ macrophages (MP), pDC initiate type I IFN-I secretion in the bone marrow (BM) of infected mice via cell-intrinsic TLR7 sensing and cell-extrinsic STING sensing. Herein we show that CD169^+^ MP and TLR7-sensing are both required for pDC arrest during priming, suggesting CD169^+^ MP are the source of TLR7 ligands. We establish that TLR7 sensing in pDC and chemotaxis are both required for pDC arrest and functional clustering with CD169^+^ MP in the BM. Lastly, we demonstrate that STING-sensing in CD169^+^ MP control pDC initiation of type I IFN production while also regulating pDC clustering and egress from the BM. Collectively, these results link pDC acquisition of type I IFN secreting capacity with changes in their motility, homing and interactions with CD169^+^ MP during infection. Thus, targeting this cellular interaction may help modulate type I IFN to improve outcomes of microbial infections and autoimmune diseases.

## Introduction

Plasmacytoid dendritic cells (pDC)(1–3) are equipped with the unique ability to rapidly secrete up to one thousand times more type I IFN -both IFNα and IFNβ- than any other cell (4–6). This is accounted for by several pDC-intrinsic and -extrinsic mechanisms. pDC-intrinsic features include the constitutive expression of the endosomal Toll-like receptors (TLR-) 7 and 9, which respectively recognize single stranded RNA and unmethylated CpG motifs containing DNA, leading to type I IFN production via MyD88 signaling and IRF7-mediated transcription. pDC also express constitutively high levels of the IFNβ-transcriptional regulator IRF7, the master regulator of their type I IFN producing capacity (7, 8). Several pDC-extrinsic mechanisms are also involved in pDC activation, during which pathogen-derived material is provided to pDC by other cells, allowing further checkpoints of pDC activation (3). However, the precise cell-cell interactions and requirements are still not well understood.

Type I interferon is a potent immunoregulatory cytokine that modulates hundreds of genes (9–11). It promotes early innate immune responses by favoring cell-autonomous defense mechanisms that rapidly restrict viral escape and growth. Type I IFN signaling induces the production of pro-inflammatory cytokines that enable the mobilization of innate immune defenses and set the stage for adaptive immune responses by enhancing antigen processing, presenting and co-stimulatory capacities of antigen presenting cells. It also directly signals to activated T and B cells, promoting expression of anti-apoptotic molecules and orchestrating their differentiation into robust effector cells. In contrast to these beneficial effects, excess or dysregulated type I IFN production that occurs in the context of auto-immune diseases, suggests that type I IFN can also promote pathology (12). While in viral infections, type I IFN generally appears as an absolute requirement for effective antiviral immune responses, in bacterial (*Mycobacterium tuberculosis, Listeria monocytogenes*) and parasitic (*Plasmodium*) infections, type I IFN can favor deleterious outcomes (13–17). This is often associated with the production of an inappropriate set of inflammatory mediators that hampers or skews the development of an effective host response and promotes damaging inflammation, as also exemplified during SARS-Cov2 infection (18).

Consistent with the described roles of type I IFN, pDC-derived type I IFN has been reported to be essential for host resistance against a range of viral infections (3, 12). It augments antiviral defense through the rapid activation of innate immune effector responses such as that of NK cells, monocytes and neutrophils (19–24). It can also modulate long-term CD8^+^ and CD4^+^ T cell survival and differentiation (25). pDC-derived type I IFN also promotes autoimmunity in SLE and dermatitis, systemic sclerosis, and type I diabetes (12, 26–28). Similarly, in a mouse model of severe lethal malaria, pDC-derived type I IFN can drive innate immune cell activation and deleterious inflammatory responses (14).

Several mechanisms are implicated in pDC initiation of type I IFN production *in vivo*. These include TLR7/9 pDC-intrinsic sensing of viral nucleic acids occurring as a result of direct pDC infection by viruses (29–32). Indirect sensing of pathogen-derived materials from non-pDC infected cells also occurs via exosomes, shedding of immature virus-containing particles or even internalization of immune complexes (22, 30, 33–36). Such sensing presents several advantages as pDC do not need to be directly infected to detect microbial pathogens as they can sense pathogen-derived nucleic acids under various forms, increasing the likelihood to detect danger signals (37). Given the very low numbers of pDC in the body, this mode of recognition further increases the likelihood of successful scanning of tissues and lymphoid organs in search of microbial pathogens while preventing potentially inappropriate activation. Collectively, these studies support the idea of pDC “cooperative sensing”, during which pDC interact with other cells or cell-derived materials, for effective pathogen sensing and subsequent initiation of type I IFN production.

In line with this concept, several studies have reported that pDC form rapid clusters upon TLR7/9-mediated activation (38–40). Systemic injection of synthetic TLR7/9 ligands or murine CMV infection induces pDC to migrate via CXCR3 and CCR7 to the marginal zone of spleens where they achieve peak production of type I IFN (38). pDC can also form stable immunological synapses with CD4^+^ T cells *in vitro* (39). At steady state *in vivo*, pDC can establish transient contacts with T cells, but after TLR9-induced activation, they make prolonged contact with T cells in the marginal zone/red pulp area. More recently, lymph node pDC behavior monitored during an ongoing viral infection showed that they were recruited via CXCR3 or CCR5, respectively either by infected subcapsular CD169^+^ MP or activated CD8^+^ T cells to the site of priming and interaction with XCR1^+^ DC (40). In this setting, type I IFN from pDC promoted either local MP-antiviral response that prevented virus spreading or more efficient antigen cross-presentation by DC to CD8^+^ T cells. Consistent with these findings, the formation of a dynamic TLR7-dependent interferogenic synapse between pDC and virus-infected cells has also been described *in vitro*, in which viral RNA and type I IFN are exchanged as a possible mechanism to potentiates pDC scanning of infected cells while providing local and targeted delivery of type I IFN (41). Thus altogether, these results suggested that tissue trafficking and localization of pDC *in situ* is essential for them to undergo activation and for their optimal function. However, the precise spatio-temporal dynamics of pDC behavior *in vivo* at steady state and during infection or inflammation, and which cells and molecular events may regulate it are poorly understood. Whether pDC initiation of type I IFN production is linked to their motile behavior *in vivo* still remains largely uncharacterized.

We have previously established that i) CD169^+^ MP prime pDC to initiate rapid type I IFN production in the BM of mice infected with lethal *Plasmodium yoelii (Py)* YM (14), ii) pDC priming involved pDC-intrinsic TLR7 and pDC-extrinsic STING sensing, iii) pDC arrested in the BM of infected mice at the time of peak type I IFN production (∼1.5 days) using intravital microscopy (IVM) while they were highly motile in the BM of naïve mice. pDC arrested near or potentially in contact with CD169^+^ MP, however, these MP are also densely distributed throughout the BM making it difficult to establish a functional interaction. CD169^+^ MP are tissue-resident MP which localize at entry ports of lymphatics in dLNs and in splenic marginal zones. They capture and sample antigens from dead cells, and act as entry points for viruses and bacteria, subsequently initiating immune defenses (42–45). Based on our prior work, here we directly test the hypothesis that pDCs need to arrest and interact with CD169^+^ MP in order to properly activate and initiate type I IFN production. We assess if MP take up *Py*-infected red blood cells (iRBC), acting as a primary source of parasite-derived materials and signals to pDCs. We also explore the molecular sensing mechanisms underlying the establishment of this functional interaction. Our results show that both CD169^+^ MP and TLR7-sensing in pDC control pDC arrest. We also establish that STING sensing in CD169^+^ MP is necessary to prime pDC production of systemic type I IFN, and to allow pDC egress from the bone marrow (BM).

## Results

### CD169^+^ macrophages take up *P. yoelii*-iRBC in the bone marrow

Since CD169^+^ MP are required to activate pDC in the BM of blood stage malaria infected mice (14), we hypothesized that CD169^+^ MP initiate pDC activation after taking up *Py*-iRBC. Flow cytometric analysis revealed the presence of two sub-populations of CD169^+^ MPs in the BM, which we defined phenotypically as F4/80^hi^ CD169^int^ (F4/80^hi^ MP subset) and F4/80^int^ CD169^hi^ (CD169^hi^ MP subset) (Figure S1A). To explore uptake of *Py*-iRBC by BM MP, we isolated RBC either from naïve (uninfected RBCs, uRBC) or *Py*-infected mice, and labeled them with the red fluorescent dye tetramethylrhodamine (TAMRA-Red^+^) before transfer to WT recipient mice for FACS and IVM analyses of the BM (Figure S1B). This strategy enabled quantification and visualization of the initial uptake phase of *Py*-iRBC versus uRBCs. For *in vivo* visualization of CD169^+^ MPs, mice were injected i.v. with anti-CD169-PE mAb 16 hrs prior to IVM imaging (14). By 6 hrs post-transfer, both F4/80^hi^ and CD169^hi^ BM MP took up significantly increased proportions of TAMRA-Red^+^ *Py*-infected iRBCs compared to uRBCs, with a ∼10-fold higher uptake in both CD169^+^ MP subsets (Figure 1A). Of note, we did not find evidence for direct uptake of *Py*-iRBCs by pDC (Figure S1C). Time-lapse IVM imaging confirmed *Py*-iRBC attachment and accumulation on CD169^+^ MPs in the BM, and possibly internalization (Figure 1B and Supplemental Movie 1). Thus, taken together these data show that both populations of CD169^+^ MP take up *Py*-iRBC in the BM of infected mice *in vivo*, consistent with the hypothesis that MPs can present parasite-derived materials and provide activating signals to pDC.

**Figure 1.**
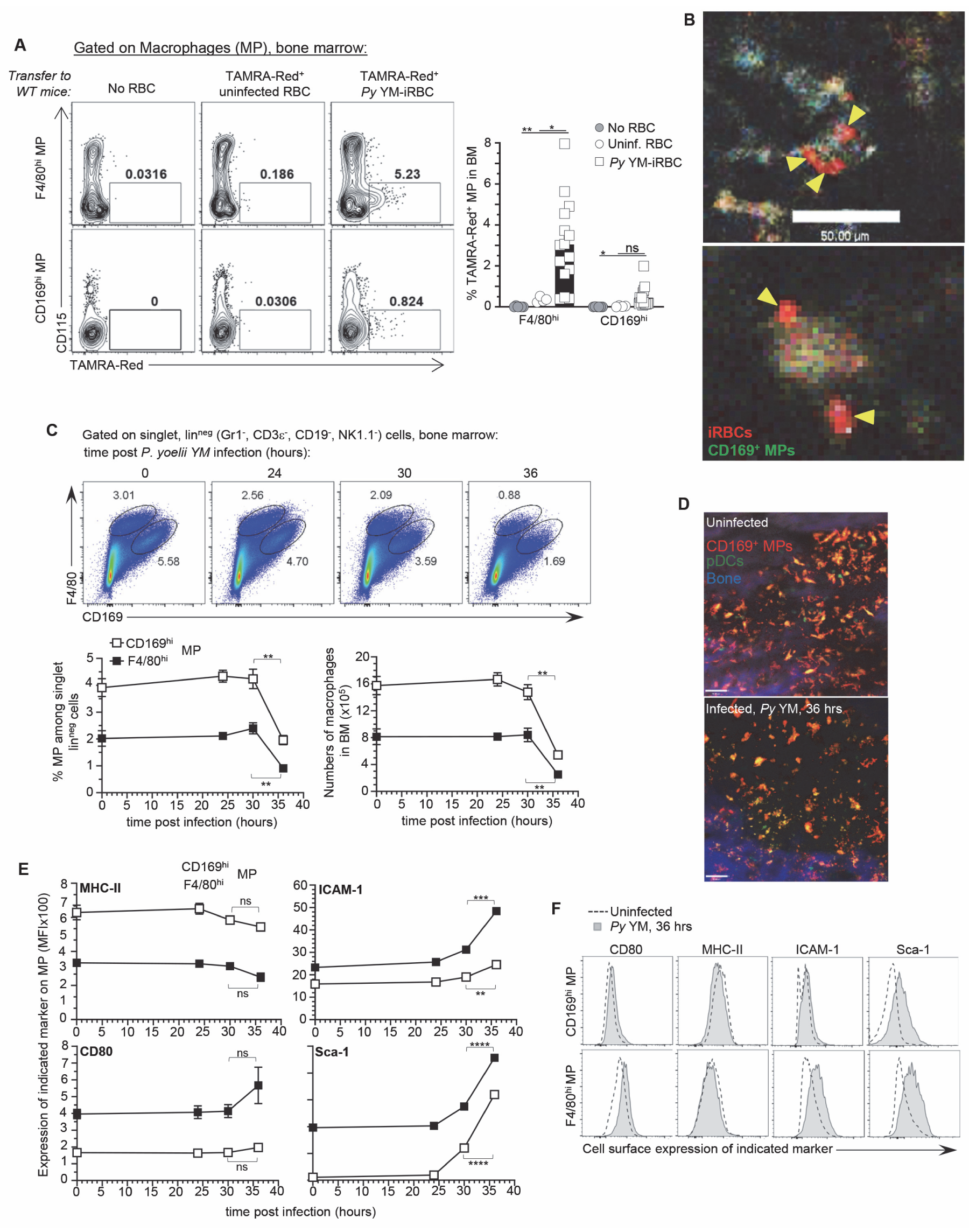
CD169^+^ macrophages uptake *Plasmodium yoelii-*infected RBC (iRBC) in the bone marrow and undergo activation and loss. F4/80^hi^ and CD169^hi^ macrophages were gated as depicted in Figure S1A. Representative dot plots of frequency of TAMRA-Red^+^ F4/80^hi^ or CD169^hi^ MP from control (no transfer), uninfected RBC (uRBC) and infected RBC (iRBC) transferred recipient mice are shown. Bar graphs show the mean frequencies +/-SEM of TAMRA-Red^+^ MP subsets in a pool of 6 independent replicate experiments with each symbol representing 1 mouse (n=4-15 mice/condition). (B) Intravital image of TAMRA-Red^+^ *Py*-iRBC (red) associated with CD169^+^ MP (green) in tibial BM 6 hr post transfer. 16 hrs prior to imaging, mice were i.v. administered CD169-FITC mAb to label CD169^+^ MP. Yellow arrowheads show *Py*-iRBC that are associated with CD169^+^ MP. Images are representative of 2 replicate experiments. (C, E, F) BM cells from uninfected or *Py* YM-infected (5x10^5^ *Py*-iRBC) WT mice were harvested at indicated time points and stained with cell-surface expression of lin^neg^ (CD3, CD19, NK1.1, CD19), F4/80, CD169 and several activation markers (MHC-II, ICAM-1, CD80, Sca-1). In (C), representative FACS dot plots of CD169^hi^ and F4/80^hi^ MP in the BM. The relative proportion and absolute numbers CD169^hi^ and F4/80^hi^ MP in mouse leg of WT B6 mice are shown in a pool of 2 replicate experiments with SEM (n=3-7 mice for each time point). (D) Representative image of tibial BM of *PTCRA-*EGFP mice administered with CD169-PE mAb 16 hrs prior to imaging either uninfected or 36 hrs post *Py*-infection in 1 of >10 imaged BM. (E, F) Representative overlay FACS histograms for indicated activation marker on CD169^hi^ and F4/80^hi^ MP from indicated mice and conditions. When relevant, *p*-values were calculated with \**p* < 0.05; \*\**p*< 0.01; \*\*\**p*< 0.001; \*\*\*\**p*< 0.0001; ns, not significant, using two-tailed unpaired Student’s *t* test.

### CD169^+^ macrophages undergo massive loss and activation in *Py*-infected bone marrow

If uptake of *Py*-iRBCs is indeed important for initial sensing of parasite-derived molecular patterns by MPs, we hypothesized that their activation status should be significantly modulated. Thus, we monitored both F4/80^hi^ and CD169^hi^ MP numbers and activation using various cell-surface markers related to type I IFN signaling (Sca-1), antigen presentation (MHC-II, CD80, CD86) and adhesion/uptake (ICAM-1, CD11a) following *Py* infection (Figures 1C-F and S1D). We found a significant loss (>50%) of both MP subsets in the BM of *Py*-infected mice, that occurred between 30 and 36 hrs post-infection as quantified by FACS analysis (Figure 1C) and IVM imaging of the tibial BM where the decrease in the overall BM MP numbers was also visualized *in situ* (Figure 1D). In addition, Sca-1, CD80 co-stimulatory molecule and ICAM-1 integrin ligand expression were increased while CD11a was diminished, consistent with robust activation of both subsets of MP in response to *Py* infection (Figures 1E, F and S1D). In summary, CD169^+^ MP in the BM of *Py*-infected mice are rapidly lost as they undergo robust activation.

### CD169^+^ MP are required for pDC arrest in *Py*-infected bone marrow

Since we previously reported that pDC arrested close to CD169^+^ MP (14), we next assessed whether *Py*-activated CD169^+^ MP control pDC arrest during infection using time-lapse IVM of the tibial marrow. As an initial step, we established baseline pDC motile behaviors using several readouts comparing uninfected versus *Py*-infected WT mice at the time pDC produce peak type I IFN (i.e., 36 hrs). We took advantage of the *PTCRA*-EGFP reporter mice, that express the enhanced green fluorescent protein (EGFP) under the control of the human pre-T cell receptor α (*PTCRA*) promoter, which is highly expressed in BM pDC (>95%, Figure S2A, (46)). Mice were also injected intravenously with a CD169-PE mAb to label BM CD169^+^ MP prior to IVM imaging. In WT uninfected mice, pDC were highly motile, exhibiting long tracks and a track velocity (TV) of 7.14 μm/min (Figure 2A, B and Supplemental Movie 2). Based on their high displacement rate (DR, 1.5 μm/min) and mean square displacement (MSD), pDC migrated through and explored a large volume of parenchymal tissue, consistent with a scanning role in the BM (Figure 2B-D). In contrast, CD169^+^ MP were sessile, whether in naïve or infected mice (Figure S2B). After *Py*-infection, pDC tracks, TV (5.13 μm/min), DR (0.93 μm/min) and MSD were all significantly reduced, consistent with their arrest. Average mean values of these distinct measurements from independent mice also reflected these findings. We next tested whether CD169^+^ MP were involved in pDC arrest after *Py*-infection. We used i) *PTCRA*-EGFP reporter mice crossed to *Cd169^DTR/DTR^* mice and ii) sublethally irradiated *Cd169^DTR/WT^* or control WT recipient mice (500 rads) reconstituted with *PTCRA*-EGFP reporter BM. These chimeras allowed to rule out any substantial impact of the DTR transgene in pDC since *PTCRA*-EGFP^+^ pDC did not express the transgene. In *Cd169^DTR^* mice, whether chimeric or not, diphtheria toxin (DT) injection induced the selective depletion of both subsets of CD169^+^ MP (Figure S2C). IVM imaging of *Py*-infected DT-treated mice revealed much longer pDC tracks compared to untreated control mice (Figure 2E). Moreover, pDC DR and MSD were significantly increased compared to untreated mice (by ∼50%, Figure 2F, G, Supplemental Movie 3). As expected, *Py*-infected, DT-treated *PTCRA*-EGFP/WT control chimeras, pDC slowed down compared to uninfected. Average mean values from independent mice were also consistent with these conclusions. Thus, these data establish that CD169^+^ MP control pDC arrest during *Py* infection and suggest that pDC arrest is linked with their ability to initiate type I IFN production.

**Figure 2.**
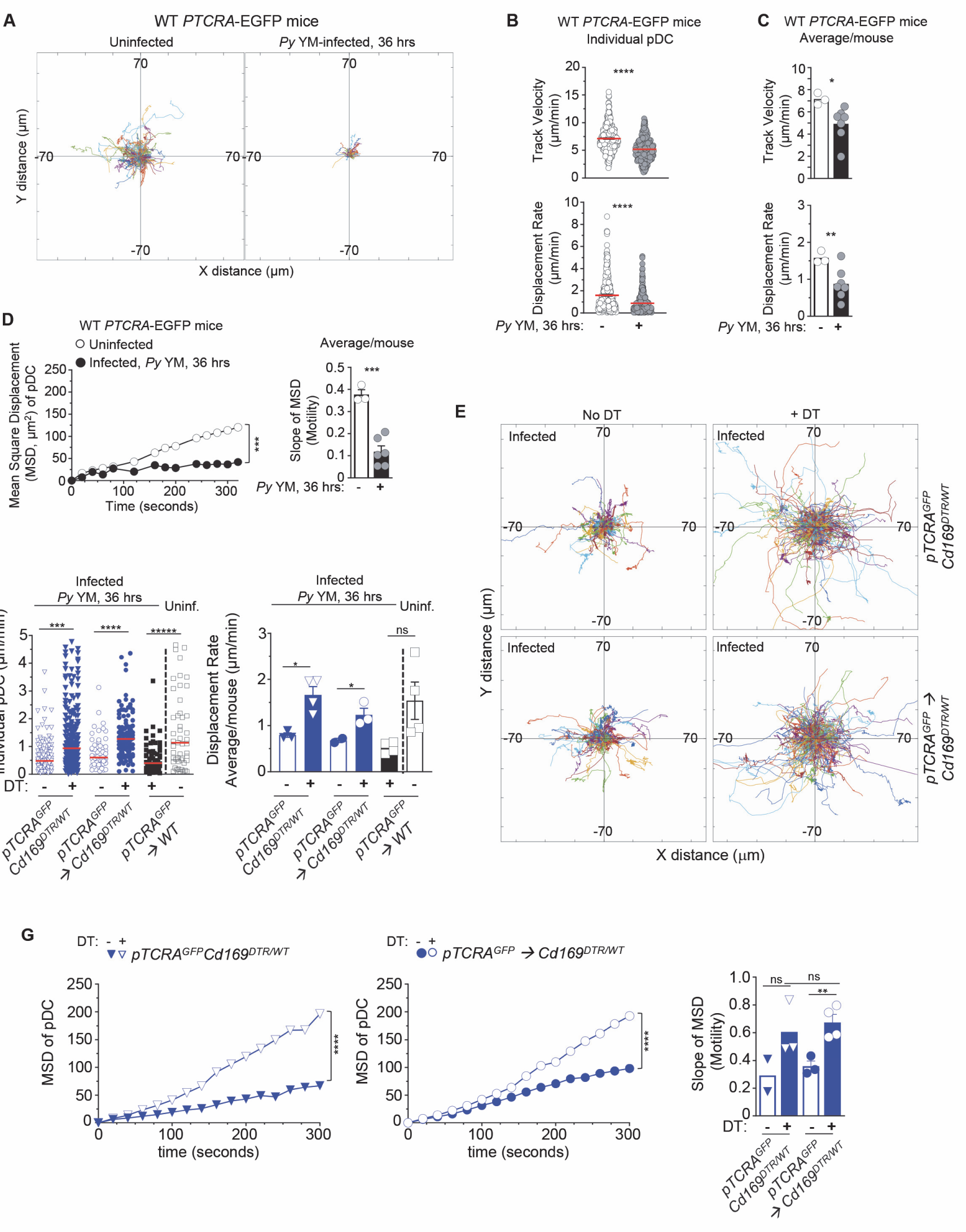
CD169^+^ macrophages control plasmacytoid dendritic cell arrest in *Py*-infected mice. *PTCRA-*EGFP reporter mice (WT, *Cd169^DTR/WT^*, or partial BM chimeras) either received PBS (uninfected) or were infected with 5x10^5^ *Py*-iRBCs, and administered with CD169-PE mAb 16 hrs prior to intravital imaging microscopy (IVM) of tibial BM. (A-D) Analysis of pDC dynamic behavior in uninfected or 36 hrs *Py*-infected WT *PTCRA-*EGFP mice. (A) Flower plots of 2D tracks of individual pDCs, superimposed after normalization of their starting coordinates to the origin. (B) Track velocity (TV) and displacement rates (DR) of individual pDC with (C) average in all mice, and (D) Mean Square Displacement (MSD) analysis of pDC over time and slope of MSD/motility. For bar graphs, each symbol features 1 mouse with mean +/-SEM (n=3-7 mice were imaged for each experimental condition and genotype). (E-G) *PTCRA-*EGFP *Cd169^DTR/WT^* or *PTCRA-*EGFP*/Cd169^DTR/WT^* and *PTCRA-*EGFP*/*WT partial BM chimeras were injected with diphtheria toxin (DT) or control PBS 12 hrs prior to *Py*-infection. (E) Tracks of individual pDC in the indicated experimental conditions. (F) Displacement rates (DR) of individual pDC and average in all mice. (G) Mean Square Displacement (MSD) analysis of pDC over time and slope of MSD/motility. For all bar graphs, each symbol features 1 mouse with mean +/-SEM (n=2-4 mice were imaged for each experimental condition and genotype). *p*-values were calculated with \**p* < 0.05; \*\**p*< 0.01; \*\*\**p*< 0.001; \*\*\*\**p*< 0.0001; ns, not significant, using two-tailed unpaired Student’s *t* test for flow-cytometry comparisons and Welch’s *t* test for comparison of IVM quantifications. Multiple linear regression analyses were applied for statistical analysis of the Mean Square Displacement Plots.

### pDC arrest in *Py*-infected bone marrow depends on TLR7-intrinsic signaling

To achieve type I IFN production, pDC require TLR7-intrinsic sensing (14). If pDC arrest is indeed linked to their optimal activation, we predicted that TLR7 sensing in pDC would also be required for their arrest. We first generated *Tlr7^-/y^* and WT *PTCRA*-EGFP reporter mice and monitored pDC motility in the tibial BM of *Py*-infected (36 hr post-infection), or control uninfected live mice using IVM imaging. In both *Tlr7^-/y^* and WT uninfected mice, pDC were highly motile with long tracks and a comparable TV (respectively 6.52 and 7.16 μm/min, Figure 3A, B and Supplemental Movie 4). However, pDC in *Tlr7^-/y^* uninfected mice migrated longer distances than in WT mice based on their track length and MSD, suggesting that tonic TLR7 signals may negatively regulate motility under steady-state conditions (Figure 3D). Following *Py*-infection in WT mice, pDC motility was reduced by 31%, in terms of TV (4.92 μm/min) and 45% based on DR (∼0.87 μm/min) as well as MSD and slope of MSD averages (Figure 3B, D and Supplemental Movie 5). In contrast, pDC in *Py*-infected *Tlr7^-/y^* mice did not arrest and continued migrating with similar speeds in terms of TV (7.05 μm/min) and DR (1.55 μm/min). (Figure 3B, C), while measurable MSD reductions were detectable (Figures 3D). Average mean values from independent mice largely confirmed these conclusions. To further assess if cell-intrinsic TLR7 sensing mediated pDC arrest, we generated partial BM chimera mice as above, in which WT recipient mice (CD45.1^+^) were grafted with either *Tlr7^-/y^*- or WT-*PTCRA*-EGFP reporter BM after sublethal irradiation. In these mice, ∼30% of hematopoietic-derived cells originated from the donor reporter BM, allowing for tracking of *Tlr7^-/y^* or control WT *PTCRA*-EGFP^+^ pDC in an environment where ∼70% of HSC-derived cells and all radioresistant cells, were intact for TLR7 signaling (Figure S3A). Consistent with the prior result (Figure 3A-D), *Tlr7^-/y^* pDC in infected chimeras exhibited long tracks and did not arrest while WT pDC had much shorter tracks and slowed down significantly, establishing that TLR7 signaling is required intrinsic to pDCfor them to stop during infection (Figure 3E, F and Supplementary Movie 6). We also quantified pDC/CD169^+^ MP contact duration as a measure of their possible interactions in chimeric mice and found that WT pDC significantly increased their contact times with the MPs during infection (from ∼2 to 16 min) whereas *Tlr7^y/-^* pDC interacted with the MP for comparable amounts of time (∼4-5 min) whether mice were infected or not (Figure 3G). Thus, taken together, these results show that TLR7 sensing in pDC is required for their arrest and interactions with CD169^+^ MP in the BM of malaria-infected mice at the time pDCs produce peak type I IFN. Since CD169^+^ MP are required for pDC arrest, it further suggests that MP are providing TLR7 ligands to pDC.

**Figure 3.**
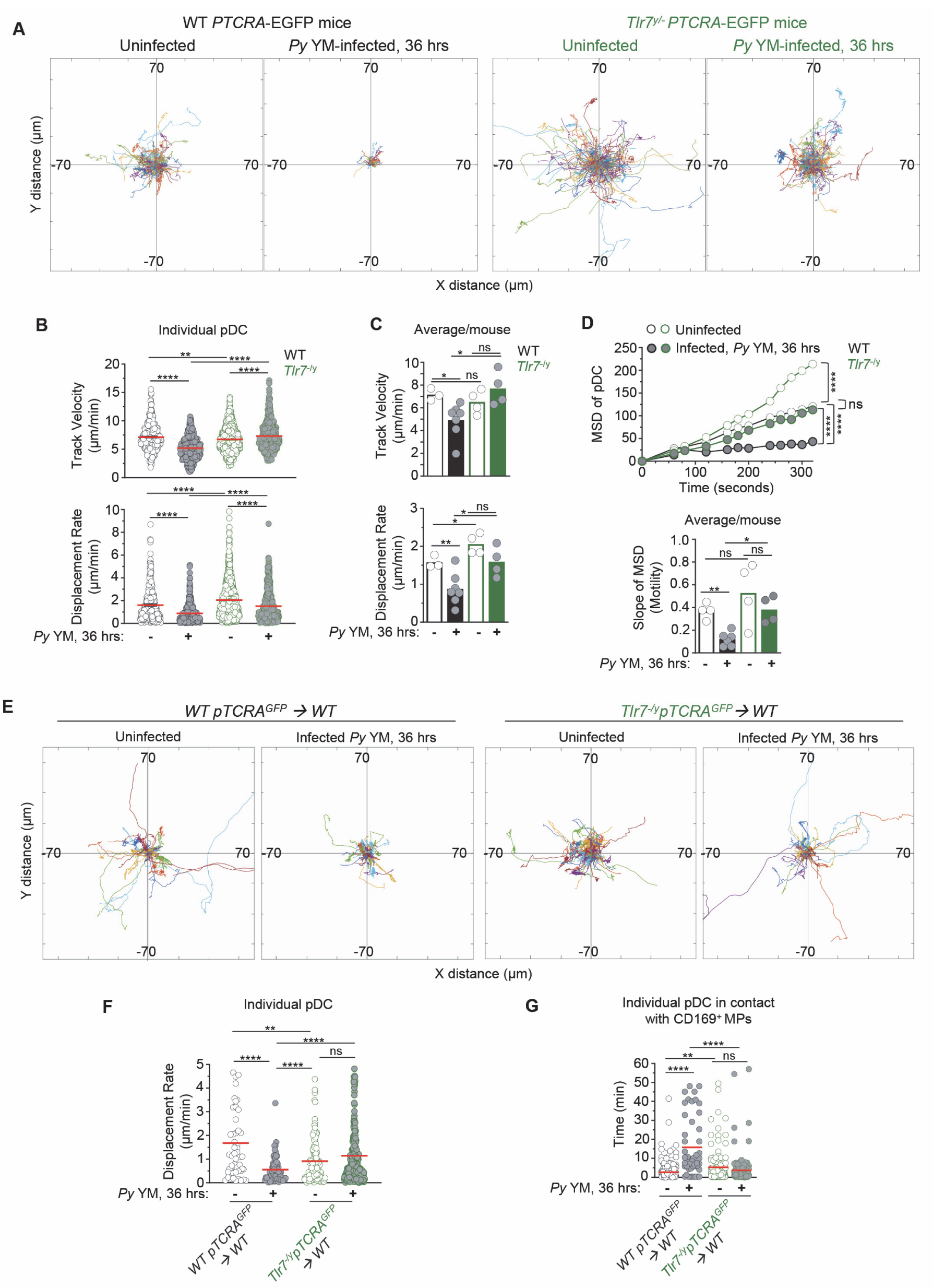
TLR7 sensing in Plasmacytoid dendritic cell is required for their arrest and clustering in *Py*-infected mice. WT and *Tlr7^-/y^ PTCRA-*EGFP reporter mice, and WT *PTCRA-*EGFP/WT and *Tlr7^-/y^ PTCRA-*EGFP/WT partial BM chimeras either received PBS (uninfected) or were infected with 5x10^5^ *Py*-iRBCs, and administered with CD169-PE mAb 16 hrs prior to IVM of tibial BM. (A) Tracks of individual pDC dynamic behavior in uninfected or 36 hrs *Py*-infected mice with starting positions in the same origin. (B) Track velocity (TV) and displacement rates (DR) of individual pDC with (C) average in all mice, and (D) Mean Square Displacement (MSD) analysis of pDC over time and slope of MSD/motility. Each symbol feature 1 mouse with mean +/- SEM. 4-6 individual mice were imaged for each experimental condition and genotype. (E) Tracks of individual pDC in uninfected or *Py*-infected WT *PTCRA-*EGFP/WT and *Tlr7^-/y^ PTCRA-*EGFP/WT partial BM chimeras. (F) DR of individual pDC and (G) quantification of individual pDC contact time with CD169^+^ MP are shown. 3-4 individual mice were imaged for each experimental condition and genotype. *p*-values were calculated with \**p* < 0.05; \*\**p*< 0.01; \*\*\**p*< 0.001; \*\*\*\**p*< 0.0001; ns, not significant, using two-tailed unpaired Student’s *t* test for flow-cytometry comparisons and Welch’s *t* test for comparison of IVM quantifications. Multiple linear regression analyses were applied for statistical analysis of the Mean Square Displacement Plots. The WT PTCRA-EGFP data depicted in: (A) are the same as in Figure 2A ; (B, C) are identical as in Figure 2B, C ; (D) are the same as in Figure 2D ; (F) are the same as in Figure 2F. This enabled comparisons across mutant mouse conditions relative to WT.

### Altered chemotactic receptor and adhesion molecule expression on pDCs from *Py*-infected *Tlr7^-/y^* compared to WT mice

Because TLR7 sensing alters pDC dynamic behavior *in vivo*, we next hypothesized that TLR7 signaling likely modulated both their migratory and adhesion characteristics.

Several chemotactic receptors (CXCR3, CXCR4, CX3CR1, CCR2, CCR5, CCR9) and adhesion molecules (CD11a, ICAM-1, VCAM-1) were reported to be expressed on pDC in naïve mice and to contribute to their trafficking and ability to cluster upon activation (38, 40, 47). We monitored cell-surface expression of chemokine receptors (CXCR3, CCR9, CXCR4, CX3CR1) and integrin ligands (ICAM-1, VCAM-1) on pDC in uninfected compared to *Py*-infected WT and *Tlr7^-/y^* mice (Figure S3). While basal expression of these receptors and integrins was comparable in all groups of uninfected mice, CCR9 and CX3CR1 were significantly upregulated on pDC after infection in WT but not in *Tlr7^-/y^* mice. Significant increases in CXCR4 expression were not reliably detectable. In contrast, CXCR3 expression was significantly downregulated on pDC in both WT and *Tlr7^-/y^* mice, but to different extents (90% and 50%, respectively). Expression of ICAM-1, a ligand for the integrin LFA-1, was significantly upregulated in both WT and *Tlr7^-/y^* after infection, but the magnitude of upregulation remained substantially greater in WT mice (factor of ∼3). Expression of CD11a, the alpha chain of LFA-1, was slightly decreased in *Py*-infected WT mice whereas expression did not change in *Tlr7^-/y^* mice. Lastly, expression of VCAM-1, ligand for the integrin VLA-4, was significantly reduced in both WT and *Tlr7^-/y^* during the infection, but to a lesser extent in *Tlr7^-/y^* mice. Collectively, these data link TLR7-mediated sensing by pDCs to changes in cell-surface expression of several chemotactic receptors and adhesion molecules that may contribute to modifications in pDC motility, homing and/or arrest that occur following *Py-*infection.

### G_α_I-protein dependent homing is required for inducing pDC production of type I IFN

Since we found important modulations in chemokine receptor expression driven by TLR7 signaling, we hypothesized that chemotaxis was involved in pDC motility and arrest. To test this possibility, we injected *Py*-infected mice with either PBS (control) or pertussis toxin (PT), which blocks G_α_I-protein coupled-receptor signal transduction and related chemotaxis, and subsequently analyzed pDC motility, morphology and type I IFN production (Figure 4 and Supplementary movie 7). In *Py-*infected *PTCRA*-EGFP reporter mice, PT-treatment led to stronger immobilization of pDC compared to control mice, with shorter tracks and significantly lower DR (Figure 4A, B). pDC arrest was also concomitant with distinct morphological changes (Figure 4C). Notably, pDCs in PT-treated mice exhibited a round shape contrasting with the elongated dendritic shape observed in PBS-treated counterparts. Quantitative analysis of individual pDC length, surface and shape factor (a measure of cell sphericity) further revealed significant differences in PT- versus PBS-treated *Py*-infected mice (Figure 4D). While after PT-treatment pDCs were immobile and unable to migrate, they were still viable in the BM and could potentially sense free diffusing parasite-derived ligands. To determine whether PT treatment impacted the ability of pDC to make type I IFN, we carried out the same experiment in *Ifnb^YFP/YFP^* reporter mice, which revealed an 80% reduction in type I IFN-expressing pDC (YFP^+^) compared to control infected group (0.44% versus 2.1% YFP^+^ pDCs, Figure 4E). Taken together with the imaging findings, this indicated that motility, chemotaxis, or any other G_α_I-protein dependent transduction process, was required to achieve maximal type I IFN-production. Despite their drastic immobilization and morphological changes, pDC in PT-treated infected mice still modulated cell-surface expression of costimulatory (CD86) and adhesion (CD62L, ICAM-1, VCAM-1) molecules, as well as chemotactic receptors (CXCR3), compared to PBS-treated infected and uninfected control mice (Figure 4E). Specifically, expression of CD62L, ICAM-1, VCAM-1 and CXCR3 but not CD86, were significantly altered in PT-compared to PBS-treated *Py*-infected mice. These results show that blocking G_α_I-protein coupled-receptor signal transduction including chemotaxis, alters expression of most but not all of these markers and the acquisition of type I IFN-secreting capacity by the pDC.

**Figure 4.**
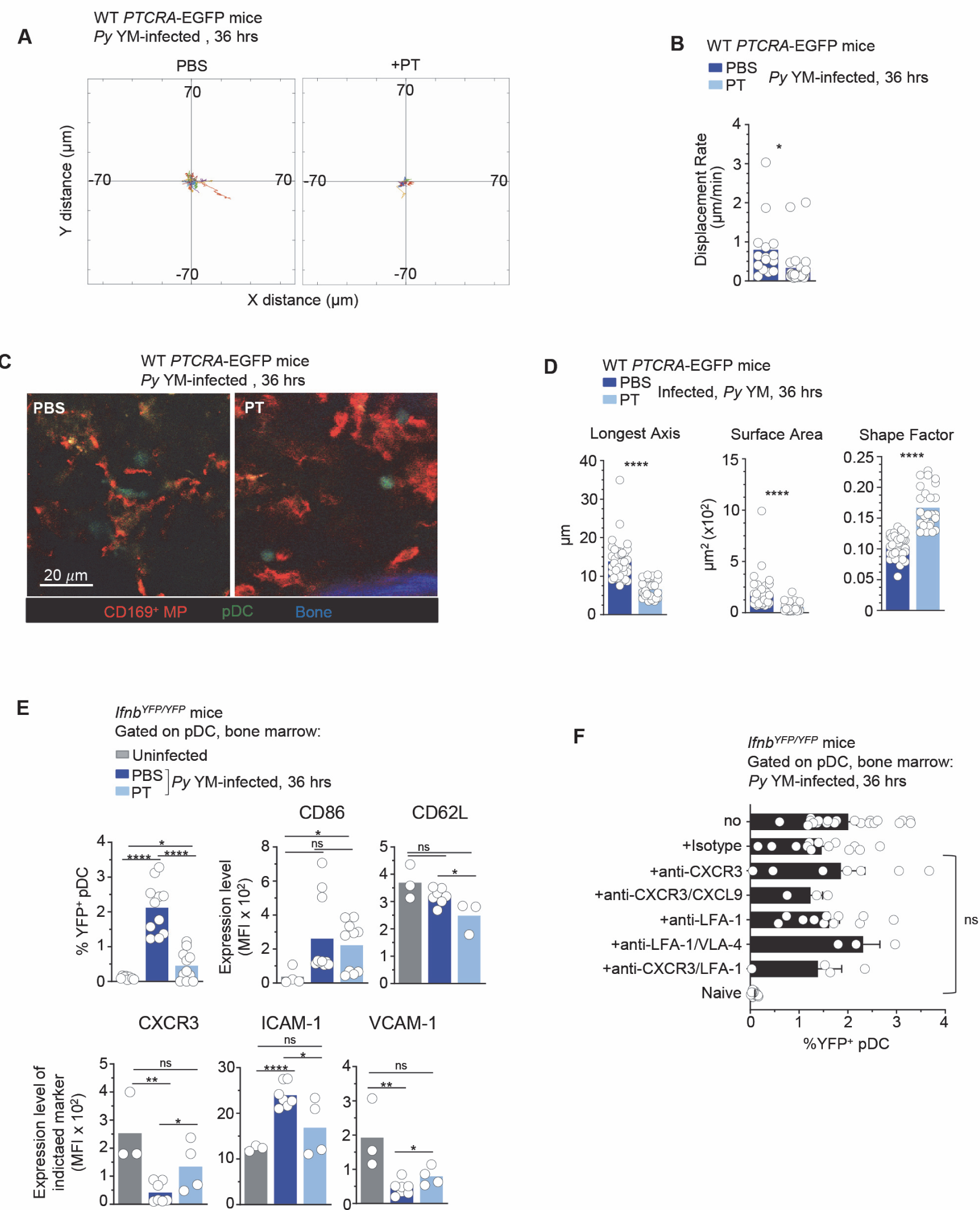
Plasmacytoid dendritic cell arrest is not sufficient to achieve robust type I IFN production during *Py* infection. WT *PTCRA-*EGFP (A-D) or *Ifnb^YFP/YFP^* (E, F) reporter mice either received PBS or pertussis toxin (PT) and were infected with 5x10^5^ *Py*-iRBCs. In (A-D), mice were injected with PT or control PBS at the time of *Py*-infection and administered with CD169-PE mAb 16 hrs prior to IVM of tibial BM. The analysis of pDC dynamic behavior of PT-treated or untreated, 36 hrs *Py*-infected mice are shown. (A) Flower plots of 2D tracks of individual pDCs, superimposed after normalization of their starting coordinates to the origin.(B) DR of individual pDC. (C) Representative IVM images of tibial BM. (D) Quantification of cell morphology of arrested pDC in PBS versus PT-treated infected mice. (E, F) *Ifnb^YFP/YFP^* reporter mice received indicated treatments (PBS, PT, Isotype Ab, anti-CXCR3+/-CXCL9+/-LFA-1, anti-LFA-1+/-VLA-4) and the time of *Py* infection, and 36 hrs later BM cells were stained for cell-surface expression of lin^neg^ (CD3, CD19, NK1.1, CD11b, Gr1), BST-2, Siglec-H, CD86, CD62L, CXCR3, VCAM-1 and ICAM-1. Bar graphs quantify the expression of YFP, CD86 and ICAM-1 by pDCs with mean+/-SEM across 2-5 independent replicate experiments, with each symbol featuring 1 mouse (n=3-10 mice). *p*-values were calculated with \**p* < 0.05; \*\**p*< 0.01; \*\*\**p*< 0.001; \*\*\*\**p*< 0.0001; ns, not significant, using two-tailed unpaired Student’s *t* test for flow-cytometry comparisons and Welch’s *t* test for comparison of IVM quantifications.

The fact that PT-treatment prevented pDC motility, and both CD169^+^ MP and TLR7-sensing were required for pDC arrest and type I IFN production, suggested that pDCs do not encounter freely circulating TLR7 ligands in the BM but rather that such ligands must be presented by *Py*-activated CD169^+^ MP through close and durable contacts with pDC. However, as PT treatment has pleiotropic activity and may induce indirect effects on other cells, we next attempted to determine more specifically which chemotactic and adhesion receptors mediated pDC homing and activation. CXCR3 is significantly downregulated after *Py* YM (Figure S3B) and has been shown to promote chemotaxis of pDC to sites of priming in the LN (40). CXCR3 has two ligands, CXCL10 and CXCL9, which are highly expressed by monocytes and MP during infections (48, 49). Thus, we tested if CXCR3 was functionally contributing to pDC activation in the BM. We injected *Ifnb^YFP/YFP^* mice at the time of infection both with anti-CXCR3 (CXCR3-173 clone) and anti-CXCL9 2A6.9.9 mAb, which blocks the CXCL10 binding site and neutralizes soluble CXCL9, respectively (Figure 4F). pDC in these mice expressed YFP at similar levels relative to control-isotype Ab-treated or non-treated mice (1.5-2%). Furthermore, Ab blocking of LFA-1 and VLA-4 integrins, with or without anti-CXCR3 mAb, also failed to decrease expression of type I IFN by the pDC, suggesting that none of the tested reagents were sufficient to recapitulate the *in vivo* effects of PT treatment on pDC activation. Collectively, these results suggested that pDC motility requires G_α_I-dependent chemotaxis. This process controls pDC homing to and/or functional interactions with CD169^+^ MP, which are essential to to sense TLR7 ligands and initiate type I IFN production.

### STING signaling is not required for pDC arrest during *Py* infection

Since i) STING signaling extrinsic to pDC is required for pDCs to secrete type I IFN during *Py* YM infection (14), and ii) pDC arrest and functional interaction with MP is linked to their full activation, we hypothesized that STING sensing could also contribute to pDC homing, arrest and interaction with MPs to initiate type I IFN production. To investigate these possibilities, we analyzed pDC motility in *Sting^Gt/Gt^ PTCRA*-EGFP reporter mice by IVM. In uninfected *Sting^Gt/Gt^* mice, most pDC exhibited high motility while they arrested in *Py-*infected counterparts (Figure 5A, Supplemental Movie 8). The majority of pDCs tracks were confined and with reduced displacement from origin following the infection. While pDC DR and MSD slopes were significantly diminished compared to naive mice, the average pDC TV remained elevated (6.52 μm/min), similar to uninfected mice (6.33 μm/min) (Figure 5B-D). Such discordance between TV and DR usually means that cells exhibit a highly confined migration pattern. Consistent with this interpretation, the MSD of pDC was significantly higher in *Sting^Gt/Gt^* versus WT infected mice, (Figure 5D). We next infected partial BM chimeras in which sublethally irradiated *Sting^Gt/Gt^* or control WT mice were reconstituted with WT *PTCRA*-EGFP donor BM, allowing to monitor WT pDC motility in a *Sting^Gt/Gt^* host where the majority of HSC-derived cells (∼70%) and all radioresistant cells are deficient for STING. In these mice where we visualized WT pDC in a *Sting^Gt/Gt^* host, pDC arrested and interacted longer with MP, as seen in control WT chimeras and in non-chimeric mice (Figure 5E, F and Supplemental Movie 9). Most interestingly, we observed that in both non-chimera and chimera *Py*-infected *Sting^Gt/Gt^* mice, pDC clusters were significantly larger in volume than in WT control mice, by 3 fold on average, and in some cases exceeding WT clusters by 10 fold (Figure 5G, H and Supplemental Movies 9, 10). We also enumerated by FACS analysis, higher numbers of pDC in the BM of *Sting^Gt/Gt^* compared to WT *Ifnb^YFP/YFP^* reporter *Py*-infected mice (Figure 5I), correlating with the significantly larger clusters of pDC. As expected, the number of *Ifnb*-expressing pDC (YFP^+^) was significantly reduced compared to WT counterparts, despite increase pDC numbers. In addition, while still significantly modulated, expression of several chemotactic receptors (CXCR3, CCR9, CXCR4, CX3CR1) and the integrin VCAM-1, failed to undergo as much cell-surface modulation as in WT mice (Figure S4). This suggested that pDC in *Sting^Gt/Gt^* mice may not receive appropriate signals to enable them to resume motility, leave clusters and egress from the BM. Taken together, these data establish that pDC-extrinsic, STING-dependent signals do not control pDC arrest, homing and interactions with CD169^+^ MP in BM during *Py*-infection, but that they may limit their accumulation and/or retention in the BM.

**Figure 5.**
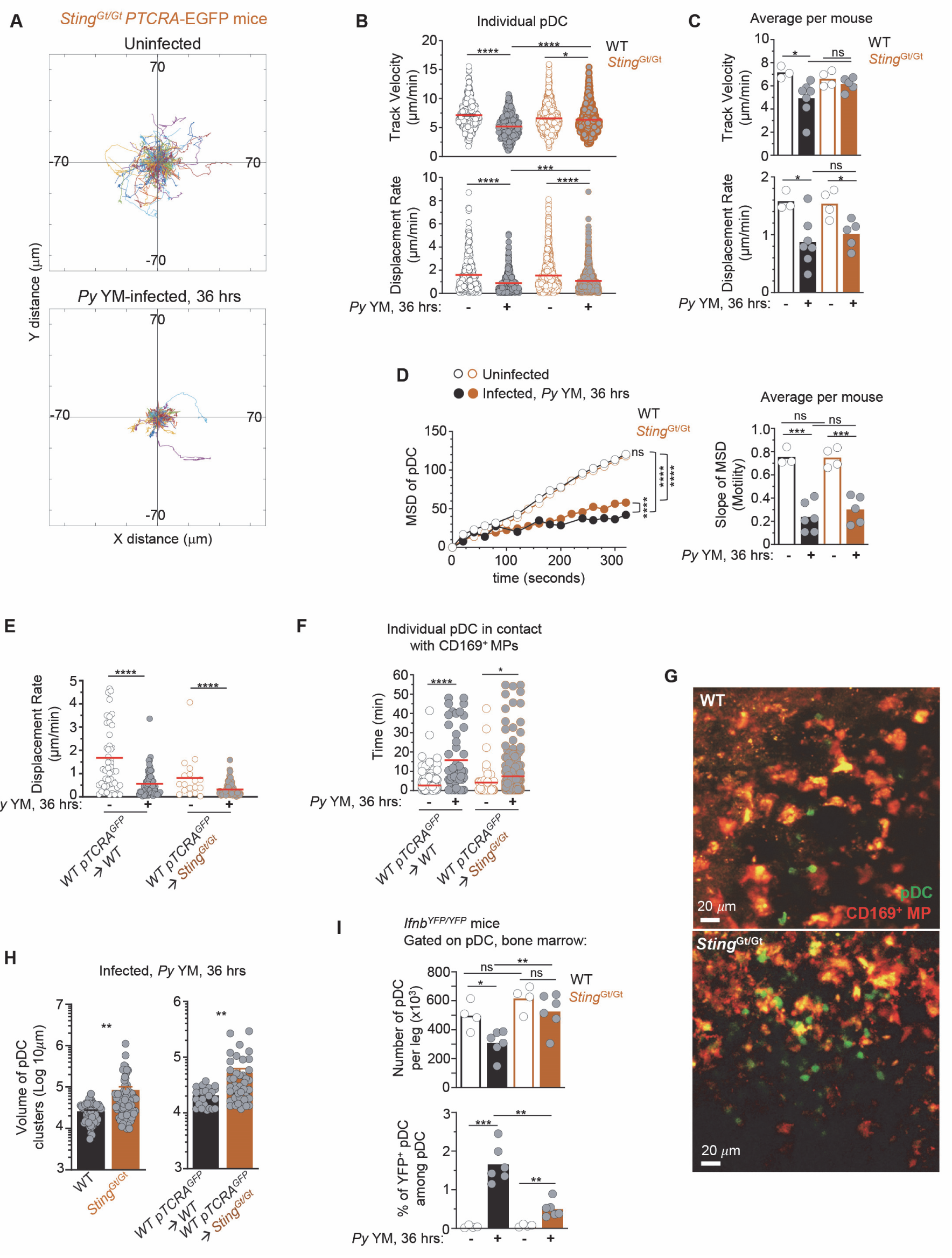
Plasmacytoid dendritic cells arrest, form larger clusters and accumulate in the BM of STING-deficient compared to WT *Py*-infected mice. WT or *Sting^Gt/Gt^ PTCRA-*EGFP reporter mice either received PBS (uninfected) or were infected with 5x10^5^ *Py*-iRBCs, and administered with CD169-PE mAb 16 hr prior to IVM imaging of tibial BM. The analysis of pDC dynamic behavior of uninfected or 36 hr *Py*-infected mice is shown. (A) Tracks of individual pDC in *Sting^Gt/Gt^ PTCRA-*EGFP reporter mice with starting positions in the same origin. (B) TV and DR of individual pDC with (C) average in all mice, and (D) MSD analysis of pDC over time and slope of MSD/motility. Each symbol features 1 mouse with mean +/- SEM. 3-7 individual mice were imaged for each experimental condition and genotype. (E) DR of individual pDC and (F) individual pDC contact time with CD169^+^ MP, in uninfected or *Py*-infected *Sting^Gt/Gt^ PTCRA-*EGFP/WT and WT *PTCRA-*EGFP/WT partial BM chimeras. (H) Quantification of cluster volume in WT, *Sting^Gt/Gt^*, *Sting^Gt/Gt^ PTCRA-*EGFP/WT and WT *PTCRA-*EGFP/WT partial BM chimeras. (G) Image from IVM of tibial BM showing pDC clusters 36 hr post *Py*-infection. For (E-H), 3-4 individual mice were imaged for each experimental condition and genotype. (I) Number of pDC per leg and relative proportion of YFP^+^ pDC in the BM of uninfected or *Py*-infected (36 hrs) WT- and *Sting^Gt/Gt^- Ifnb^YFP/YFP^* reporter mice, as quantified by FACS after staining for cell-surface expression of lin^neg^ (CD3, CD19, NK1.1, CD11b, Gr1), BST-2, Siglec-H. Bar graphs quantify the expression of YFP in pDCs with mean +/- SEM across 2 independent replicate experiments (n=4-6 mice/group). *p*-values were calculated with \**p* < 0.05; \*\**p*< 0.01; \*\*\**p*< 0.001; \*\*\*\**p*< 0.0001; ns, not significant, using two-tailed unpaired Student’s *t* test for flow-cytometry comparisons and Welch’s *t* test for comparison of IVM quantifications. Multiple linear regression analyses were applied for statistical analysis of the Mean Square Displacement Plots. Data of WT *PTCRA-*EGFP depicted in: (B, C) are the same as in Figures 2B,C and 3B, C ; (D) are also in Figure 2D, 3D. WT *PTCRA-*EGFP/WT partial BM chimera in (E) are also in figures 2F, 3F. These are included for ease of comparisons across conditions relative to WT.

### STING signaling in CD169^+^ macrophages controls pDC production of type I IFN and egress from the bone marrow

Next, we assessed if STING signaling specifically in CD169^+^ MP during *Py* YM infection controlled i) initiation of type I IFN production by pDC, and ii) pDC accumulation versus egress from the BM. We bred *Cd169^Cre/Cre^* knock-in mice to *Sting^Gt/Gt^* or *Sting^F/F^* (where exons 3-5 of *Sting* were flanked with two LoxP sites) mice to generate *Cd169^Cre/WT^Sting^Gt/F^* mice in which CD169^+^ MP lacked expression of functional STING. All these mice were also homozygous for the *Ifnb^YFP^* reporter transgene, allowing us to monitor type I IFN expression by FACS. Upon *Py* YM infection, while ∼2.2% of pDC (∼4,200 pDC/leg) in WT and *Sting^Gt/WT^* control mice expressed IFNβ (YFP^+^ ICAM-1^hi^), we quantified a ∼60% reduction in YFP^+^ pDC when STING was lacking in CD169^+^ MPs (Figure 6A), demonstrating that STING signaling in CD169^+^ MP was required for pDC priming of type I IFN producing capacity. Other markers strongly modulated on pDC upon activation (ICAM-1, CXCR3, CD86) underwent similar changes as in WT mice, suggesting multistep activation or independent processes (Figure S6A). Interestingly, in these mice where CD169^+^ MP lacked STING, the number of pDC in BM was ∼1.5 times higher than in WT or *Sting^Gt/WT^* control groups (p-value=0.1, Figure 6B). Similar increases in numbers of pDC were also enumerated in the BM of whole-body STING-deficient (*Sting^Gt/Gt^*) or CD169^+^ MP-depleted (DT-treated *Cd169^DTR/WT^*) infected mice, consistent with retention of pDC in BM when CD169^+^ MP or STING-dependent signals were absent (Figure 5G-I and Supplemental movies 9, 10). Collectively, these results show that STING signaling in CD169^+^ MP during *Py* infection, provide pDC with key signal(s) to initiate type I IFN production and egress from the BM.

**Figure 6.**
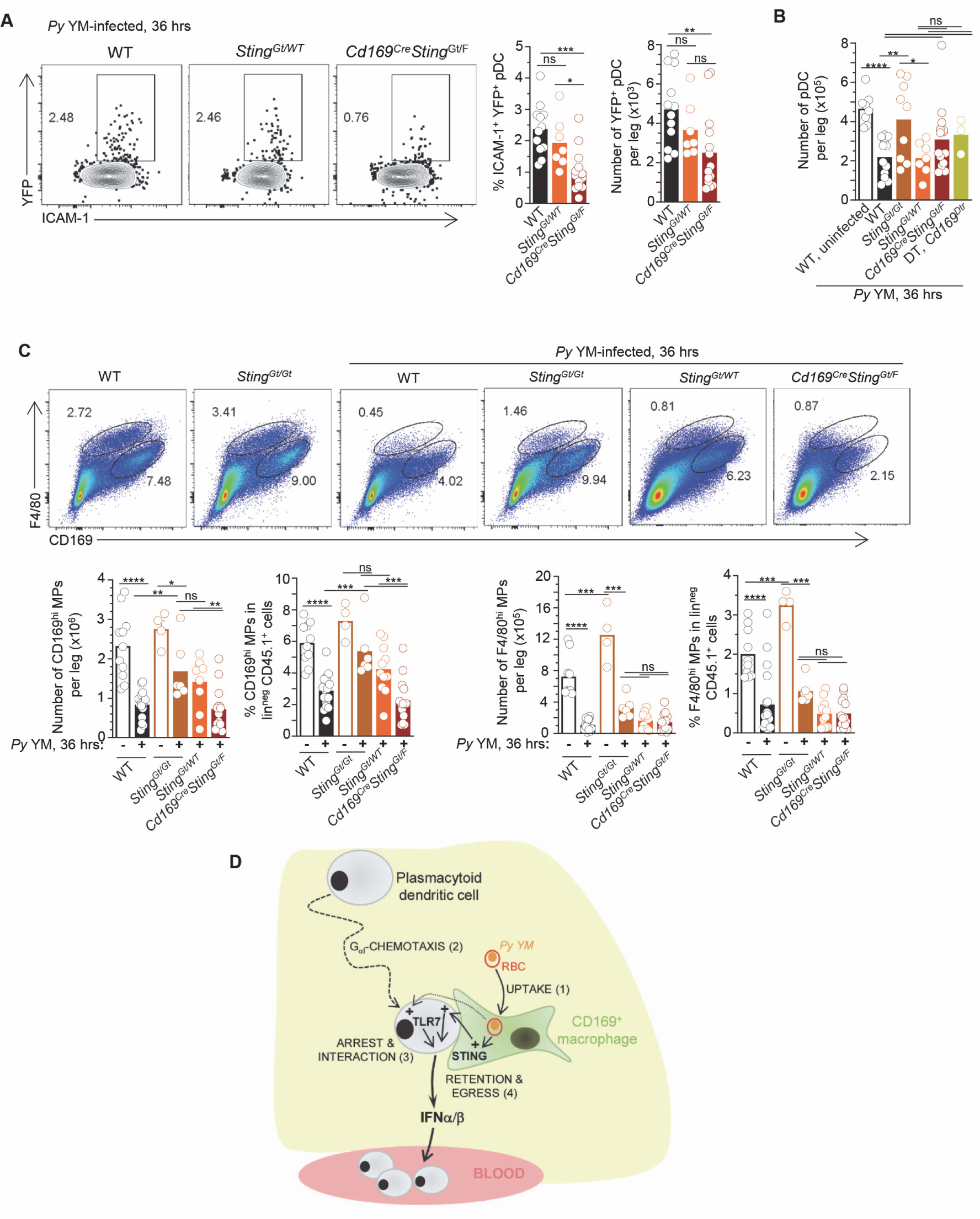
STING-signaling in CD169^+^ macrophages is required for plasmacytoid dendritic cells production of type I IFN and egress from the BM. WT-, *Sting*^Gt/WT^- or *CD169^Cre/Cre^Sting^Gt/F^*-*Ifnb^YFP/YFP^* reporter mice were infected with 5x10^5^ *Py*-iRBCs. 36 hr later BM cells were stained with live/dead and for cell-surface expression of lin^neg^ (CD3, CD19, NK1.1, Gr1), CD11b, BST-2, Siglec-H, CD169, F4/80 and indicated cell surface markers. (A) Representative FACS dot plots of YFP/ICAM-1^+^ pDC are shown, together with the proportion and numbers of YFP^+^ pDC/leg in summary bar graphs. (B) Number of pDC per leg in indicated mouse genotype. (C) Representative dot plots of CD169^hi^ and F4/80^hi^ MP in indicated mice. Bar graphs show numbers per leg and proportions. Data represent the pool of 3 independent replicate experiments with each symbol featuring 1 mouse (n=3-18 mice). *p*-values were calculated with \**p* < 0.05; \*\**p*< 0.01; \*\*\**p*< 0.001; \*\*\*\**p*< 0.0001; ns, not significant, using two-tailed unpaired Student’s *t* test. (D) Working model: during *Py* YM malaria infection, CD169^+^ MP rapidly take-up *Py*-infected RBC (step 1). Next, pDC, which are highly motile in the BM of naïve mice, home to and establish interactions with CD169^+^ MP at peak production of type I IFN by pDC (steps 2&3). This process is dependent on G_aI_-mediated chemotaxis. CD169^+^ MP and TLR7 sensing in pDC are both required for pDC arrest. Lastly, STING signaling in CD169^+^ MP is needed for pDC to initiate type I IFN secretion, regulate pDC clustering/retention, and egress from the BM to the blood (step 4).

### STING-mediated activation of pDC is independent of CD169^+^ MP loss

Upon *Py* infection, a majority of CD169^+^ MP are rapidly lost (Figure 1). We hypothesized that STING-signaling may control their loss and thus contribute to providing activating ligands and signals to pDC. We therefore assessed whether STING contributed to MP loss during *Py* infection. In whole-body *Sting^Gt/Gt^* mice, CD169^hi^ but not F4/80^hi^ MP, were partially protected from loss compared to WT control mice (by ∼50%, Figure 6C), suggesting that this may prevent optimal pDC priming. However, in *Cd169^Cre/WT^Sting^Gt/F^* mice where STING is selectively lacking in CD169^+^ MP, both MP populations underwent similar loss as in WT mice, suggesting STING deficiency extrinsic of MP protects CD169^hi^ MP from loss. However, in *Cd169^Cre/WT^Sting^Gt/F^* mice, pDC failed to initiate type I IFN transcription and accumulated in the BM compared to control groups (Figures 5I and 6B), consistent with a model in which STING signaling in CD169^+^ MP provide activating signals to interacting pDC that are unrelated to their loss. Of note, MP activation 36 hrs post infection, as measured by cell surface modulation of MHC-II, CD80, ICAM-1 and Sca-1, was mostly comparable in all infected groups, likely ruling out a general activation defect of STING-deficient MP (Figure S6B). In summary, these data uncouple MP loss that occur during *Py* infection from STING-mediated priming of pDC by CD169^+^ MP.

## Discussion

Here we reveal a complex multi-cell choreography that orchestrate pDC production of type I IFN during malaria infection. We establish the existence of a functional interaction between CD169^+^ MP and pDC *in vivo* that is needed to license pDC priming of type I IFN production *in vivo* during this infection. We link pDC arrest to their optimal priming. We show that CD169^+^ MP take up parasite-infected RBCs in the BM. We find that both CD169^+^ MP and pDC-intrinsic TLR7 sensing are required for pDC arrest, suggesting that MP are providing TLR7 ligands to pDC. We also show that MP-intrinsic STING sensing is required to activate pDC production of type I IFN, and enable their egress from the BM after priming (Figure 6D).

These findings are therefore consistent with a model in which CD169^+^ MP in the BM act as first line responders to infection, akin to other CD169^+^ MP found in the LN and spleen (50). Depletion of these MP prior to infection prevents type I production by pDCs (14), indicating that these APCs are non-redundant and are uniquely capable of presenting *Py* ligands and/or other innate signals to pDC in the BM. The fact that depletion of CD169^+^ MP prevents pDC arrest, but that forcing pDC arrest by blocking chemotaxis failed not rescue their full activation, establishes that pDC motility, homing and interaction with MP are prerequisites to achieve type I IFN production. These results suggest that TLR7-ligands, which trigger pDC arrest and type I production, are not freely diffusing in the BM and/or that *in vivo,* pDC require free nucleic acids to be properly presented or packaged to access TLR7-containing endosomes in pDC. In either case, we propose that interactions with CD169^+^ MPs are required to deliver *Py*-derived ligands to pDCs in the BM. This finding is consistent with prior studies showing pDC sensing of hepatitis C infected hepatocytes via viral RNA-containing exosomes (33, 34) or cellular particles containing Dengue virus (30). pDC were also reported to be directly infected by specific viruses such as herpes simplex virus and arenaviruses, and to utilize specialized intracellular processes like autophagy to sense viral nucleic acids and initiate type I IFN production (29, 31, 41). Given pDC very low frequency and numbers and their rapid response, it is likely that pDC are largely using such indirect but more effective modes of recognition. The formation of specialized cell-cell structures between pDC and infected cells or cells presenting pathogen-derived molecules, is indeed proposed to mediate both uptake and sensing of TLR ligands and the local delivery of type I IFN in interferogenic synapses (41).

Proper clustering and arrest of pDCs is a TLR7 guided process. TLR7 is required in pDC for their arrest, and type I production, as well as extensive remodeling of pDC expression of chemokine receptors and adhesion molecules, likely promoting stable contacts with activated, *Py*-loaded MPs. *In vivo*, pDC are recruited via CXCR3- and CCR5-dependent chemotaxis to virus-infected subcapsular draining LN MP or CD8^+^ T cells, respectively, for local delivery of type I IFN, rapidly restricting early virus spreading while promoting effective T cell priming (40). Given that pDC express multiple chemotactic receptors, and some are significantly up or downregulated during *Py* infection (e.g., CCR9, CXCR3, CX3CR1), these processes are likely to be redundant. Consistent with this interpretation, blockade of CXCR3-mediated chemotaxis failed to recapitulate all G_α_I-mediated chemotaxis blockade. Even co-blockade of several adhesion molecules (LFA-1, VLA-4) involved in classical leukocyte adhesion and clustering, did not abrogate the production of type I IFN by pDC, suggesting that other chemotactic and adhesion mechanisms are involved and need further examination.

Finally, we establish TLR7-dependent arrest and interaction of pDC with CD169^+^ MP. We also show that full activation of pDC requires STING signaling in CD169^+^ MP and that MP control pDC arrest. It was therefore unexpected that we did not find STING to control pDC arrest. However, we revealed that in STING-deficient mice, pDC formed significantly greater size clusters and accumulated in the BM compared to WT counterparts. This result was confirmed in mice lacking STING in the radioresistant compartment and only in CD169^+^ MP, consistent with a model in which STING-dependent signals control pDC egress from the BM, and as a consequence, the organization of the clusters. In our previous work (14), we reported that pDC numbers in the BM of *Py*-infected mice decreased after peak type I IFN production while that of blood pDC increased. Larger clusters and increase numbers of pDC in STING-deficient mice, and in mice with STING-deficient CD169^+^ MP, may indicate that pDC are not fully activating and thus remain docked, awaiting for additional signals. These large clusters of pDC are further evidence that pDC priming is spatially-segregated in the BM and that activation signals are not freely diffusing. These results further connect the choreography of the cell-cell signal transmissions with pDC changes in arrest and migration for efficient regulation. Other signals that need to be elucidated are likely to regulate these cellular interactions *in vivo*.

In particular, which STING-dependent signal(s) are delivered from *Py*-activated CD169^+^ MP to pDC remains unknown. STING was originally identified as a cytosolic nucleic-acid sensor, which triggering leads to the induction of a robust type I IFN response via IRF3 and NFkB, and a rapid antiviral state that enables the control of viral infections (51, 52). While pDC can secrete type I IFN independently of type I IFN signaling during VSV and MCMV infections (53, 54), pDC are also known to require type I IFN autocrine signaling when responding to TLR7/9 synthetic ligands *in vivo* (38). Thus, type I IFN may be the missing STING-dependent signal during malaria infection. Since STING also triggers the proinflammatory transcriptional regulator NFkB, other cytokines could be the missing signal. STING signaling can also trigger distinct inflammatory cell deaths and autophagy as a defense mechanism. Among STING-mediated cell death, lysosomal cell death, pyroptosis, necroptosis and NETosis were described to occur in myeloid cells in response to cGAS-STING sensing/signaling of viral DNA (55–58). While we observed a major loss of CD169^+^ MP in the BM (>60%) of *Py*-infected WT mice when type I IFN levels peak (30-36 hrs), STING-deficient MP underwent similar loss, likely excluding STING involvement in MP death. However, STING-driven autophagy, could provide live CD169^+^ MP with *Py*-derived nucleic acids and other products for optimal presentation to pDC. MP are required for pDC to arrest and initiate type I IFN-production, however, the precise sequence of events still remains unclear. Presumably, selective elimination of CD169^+^ MPs using the CD169 depleter mouse model occurs within 12 hr post DT injection, which prevents pDC production of type I IFN. The massive loss of MPs during the natural course of *Py* infection occurs after 30 hrs, suggesting that this partial loss of MP is a key process contributing to optimal pDC activation. Blocking pDC chemotaxis within a few hr post-infection, also prevents pDC to undergo full activation. Taken together, these data support a model in which pDC need to migrate in apposition to CD169^+^ MP that have acquired *Py*-derived products to establish functional interactions leading to type I IFN secretion.

STING is generally shown to require cGAS for cytosolic double stranded DNA recognition (15) and the production of STING-activating cyclic dinucleotides second messengers (59). Our results are discrepant with a prior study that also used lethal murine *Py* YM, and concluded that STING-sensing of *Py* parasites prevented strong type I IFN responses by pDC through SOCS1 inhibition of TLR7/MyD88 signaling in pDC (60). In our previous work, as well as that of others (14, 15), blocking type I IFN production and signaling, increased mouse resistance to severe *Py* and *P. berghei* murine malaria, respectively. We neither noted increased type I IFN levels in *Sting^Gt/Gt^* mice, nor increased IFNβ production by STING-deficient pDC, and *Sting^Gt/Gt^* mice did not exhibit greater resistance to *Py* YM either. While the STING pathway may be active in pDCs, current experimental evidence are scarce (3, 61), consistent with our finding that STING sensing occurs in the CD169^+^ MP, extrinsically to pDCs. Taken together, we favor the interpretation that the *Py* YM strain used in this previous work is likely to exhibit genetic differences with the repository *Py* YM strain MRA-755 we have used, which may account for the strikingly different outcomes of the two studies.

In conclusion, our analysis of the dynamic behavior of pDC *in vivo* by IVM directly in the BM of mice undergoing severe murine malaria, reinforce a model in which pDC sensing of their ligands *in vivo* is a highly coordinated and orchestrated process. We establish that CD169^+^ MPs, a subset of tissue-resident MP that act as essential sentinels of the immune system constantly sampling the lymph and blood (42–45), play a key role in regulating pDC activation *in vivo*. While we establish the existence of a functional interaction between these MP and the pDC in the BM of malaria-infected mice, further investigations will be needed to determine i) the nature of the STING-dependent signals derived from CD169^+^ MP that allow pDC to achieve type I IFN-producing capacity and egress the BM to the blood and ii) whether this interaction also takes place in other tissues and infections or autoimmune diseases.

### Materials and Methods Mice

C57BL/6J wild-type (WT B6), *Tlr7^y/-^* (62) (strain 8380, Jackson Labs), *Ifnb^mob/mob^/Ifnb-yfp^+/+^* (denoted *Ifnb*^YFP/YFP^)(63) (strain 10818, Jackson Labs), *Sting^Gt/Gt^* (64) (strain 17537, Jackson Labs), *Sting^F/F^* (strain 031670, Jackson Labs) mice have been described and were purchased from the Jackson laboratory. *PTCRA*-EGFP mice (129 background)(46) and were obtained from Dr. Boris Reizis (NYU, New York, NY) and backcrossed to WT B6 mice for 10 generations. *Tlr7^-/y^* and *Sting^Gt/Gt^ PTCRA-*GFP as well as *TLR7^-/y^* and *Sting^Gt/Gt^*, *Ifnb^YFP/YFP^* reporter mice were obtained by intercross respectively for use in intravital microscopy and FACS assays. *Cd169^DTR/DTR^* and *Cd169^Cre/Cre^* mice, a kind gift from Dr. Masato Tanaka (Riken, Japan) (45) expressed the human diphtheria toxin receptor (DTR) or the Cre recombinase knocked-in to the exon 1 of the CD169-encoding gene the mice. Both mice were used as heterozygous for experiments (*Cd169^DTR/WT^* and *Cd169^Cre/WT^Cd169^Cre/WT^Sting^Gt/F^Ifnb^YFP/YFP^* or *Sting^Gt/F^Ifnb^YFP/YFP^* were obtained by intercrossing of the relevant strains above. All mice were housed and bred in specific pathogen-free conditions at the Albert Einstein College of Medicine. 8-12 weeks old and sex-matched males and female mice were used for all experiments.

### Partial bone marrow chimeras

*Cd169^DTR/WT^* or WT B6 recipient mice were exposed to 5 Gy total body irradiation (500 rads) and reconstituted with 2×10^6^ BM cells isolated from either *Tlr7^y/-^* or WT *PTCRA*-EGFP reporter mice. Chimerism of reconstituted mice was checked ∼6 weeks later in the blood, before infections were conducted. Blood chimerism of donor-derived HSC was ∼30%.

### Ethic Statement

Animal Research: This study was carried out in strict accordance with the recommendations by the animal use committee at the Albert Einstein College of Medicine. The institution is accredited by the “American Association for the Use of Laboratory Animals" (DHEW Publication No. (NIH) 78-23, Revised 1978), and accepts as mandatory the NIH "Principles for the Use of Animals". All efforts were made to minimize suffering and provide humane treatment to the animals included in the study.

### Plasmodium infections

*Plasmodium yoelii yoelii (17X), clone* YM parasites (stock MRA-755) was obtained from BEI Resources. *P. yoelii* YM-infected red blood cells (iRBCs) from a frozen stock (stored in liquid nitrogen in Alsever’s solution and 10% glycerol) were injected intraperitoneally (i.p.) into a 8-12 week old WT B6 mouse, and grown for ∼4 days. When parasitemia reached 2–10%, 5x10^5^ *P. yoelii* YM iRBCs were inoculated intravenously (i.v.) in a volume of 200 μl PBS into each experimental mouse.

### Preparation of cell suspensions for flow cytometry (FACS) analysis

Blood was collected in K2 EDTA tubes and red blood cells lysed with NH_4_Cl buffer (0.83% vol/vol). Bone marrow was harvested from the femur and tibia by removing epiphyses to expose marrow cavity and placing into a 0.5 mL Eppendorf with a punched hole in the bottom made with an 18.5G needle, nested into a 1.5 mL Eppendorf tube. Tubes were next centrifuged at 10,000xg, 4°C, and cells resuspended in media for further used for further analyses (see below).

### Cell staining for FACS analysis

Cell suspensions were stained with LIVE/DEAD Fixable Aqua (Thermo Fisher Scientific) or Ghost Dye Red 780 (Tonbo) in PBS for 30 min at 4°C, then incubated with Fc-block CD16/32 (clone 2.4G2) for 20 minutes at 4°C. Cell suspensions were next incubated in Ab mixes in FACS buffer (2 mM EDTA, 2% FBS, 0.02% sodium azide in PBS) for 20 min at 4°C. For cell counts, we utilized 50,000 counting beads (SpheroTech) added to cells isolated from one leg or one spleen. Stained cells were collected either on a FACS BD LSR-II or Aria III. A list of all mAbs and sera used in the study with full information is provided in **Table S1**. Data were analyzed using FlowJo version 9.6.2 (TriStar).

### RBC labeling

Red blood cells were harvested from uninfected or *Py* YM-infected WT B6 mice with ∼20% parasitemia by submandibular puncture into K2 EDTA tubes and washed twice with 10mL 1X Mg^2+^Ca2^+^-free PBS. Peripheral blood was pelleted by centrifugation at 800xg and 4°C for 5 min. RBCs were gently resuspended up to a volume of 5 mL in 1X Mg^2+^Ca2^+^-free PBS, and overlaid on top of 5 mL room temperature Ficoll-Plus in a 15 mL falcon tube which was centrifuged at 1,000xg for 15 min at 20°C without breaks.

RBC pellet was washed in 5 mL Mg^2+^Ca2^+^-free DPBS, before staining with 5μM 5-Carboxytetramethylrhodamine (TAMRA-Red, ThermoFisher Scientific, Catalog#C6121) in PBS 2%FBS for 20 min at 37°C, 5%CO2 and in the dark. Labelled RBCs were then washed in 10 mL PBS, and 50μL of packed RBCs recovered in 200μL PBS were injected i.v. by retro-orbital route into recipient mice.

### Pertussis Toxin treatment

Pertussis toxin (PT, Sigma Millipore #516560) was reconstituted in 100mM sodium phosphate, 500 mM sodium chloride, pH 7.0 and stored at -80°C. Mice were treated with 2.5µg/µl pertussis toxin in 100µl of 1xPBS, i.v. at the time of infection.

### Two-photon preparation, imaging and analysis

*In vivo* imaging of BM was conducted as described previously (65). *PTCRA*-EGFP reporter mice were anesthetized with isoflurane (2.5-3% for induction, 1-2% for surgery/imaging) admixed in 1:1 O_2_: air mixture at a flow rate of 1L/min. Once induction was achieved, a gas mask was secured over the nose and stabilized using Velcro on a stage maintained at 37°C. The mouse anesthetic plane was ensured through the lack of response to toe and/or tail pinches. Level of isoflurane was adjusted accordantly to ensure the mouse maintained appropriate sedation with a stead, non-labored respiratory rate between 60-80 breath/min. Leg hair was then shaved using a surgical blade and a small amount of soap and water to facilitate hair removal. Loose hair and residual soap/water was removed with a kimwipe and the leg dried. Subsequently, the mouse torso was secured on the stage with Velcro under the ribcage so as to not interfere with breathing, and the leg stabilized with a piece of scotch tape with the mouse is a supine position and the foot everted to reveal the medial surface of the leg. Then, the medial aspect of the tibia was surgically exposed by making a small incision at the level of the medial malleolus, avoiding the saphenous artery, to expose the tibial bone. A window was secured to the leg using vet bond, sliding a small tooth of the window under the tibial bone and then fixed to the stage using screws. Then, the exposed bone was shaved using a microdrill to a 100μ thickness. Once optimal thinness was achieved, a ring of vacuum grease was drawn around the imaging area, then filled with a drop of 37°C lactated ringers (Thermo Scientific Cat#R08432), and the mouse transferred to the microscope chamber maintained at 37°C. Images were collected on an Olympus FV-1000MPE upright laser scanning microscope using a MaiTai DeepSee Laser (Spectraphysics) set to a wavelength of 910 or 920 nm. BM was visualized using a 25X NA 1.05 objective and magnified 1.5-2x. Time lapse stacks with 5 μ slices spanning a total depth of 50 μ were captured at an acquisition of 4.0 μs/pixel using. To visualize CD169^+^ MP, mice were intravenously injected with 5μg of anti-CD169 (SER4) antibody conjugated to PE (BD Pharmingen), 16 hrs prior to imaging (66).

#### Movie Analysis

4D image sets were analyzed on Volocity 6.3 (Improvision). For cell-contact duration between pDC and CD169^+^ MP, analysis was conducted by tallying time durations for all pDC in the field in contact with CD169^+^ MP. Data was pooled from 3 or more mice from independent experiments. Supplementary movies were assembled using After Effects (Adobe). Contrast and saturation were enhanced to allow for greater ease of pDC visualization above MP.

Two-dimensional plots were created at a higher resolution to our specification using Matlab. (Instructions and script hyperlinked: <u>../XYZ plots/Read Me.docx</u> ; <u>../XYZ</u> <u>plots/XY_Plots_ZB_10082019.m</u>

### Statistics

All statistical tests were run with GraphPad Prism 8. Significance is depicted as follows: *p<0.05; **p<0.01; ***p<0.001; ****p<0.0001. All graphs show mean +/- SEM. The unpaired *t* test was used for comparing 2 groups related to FACS data. Welch’s *t* test was used for comparing 2 groups related to IVM analyses. Multiple linear regression analyses were applied to test if the factor conditions (time and group) predicted displacement for statistical analysis of the Mean Square Displacement Plots, assuming a gaussian distribution of residual and least squares.

## Acknowledgements

We thank the AECOM FACS core facilities. We thank Ilseyar Akhmetzyanova for helping with initial training on intravital microscopy and Zachary Benet for providing the MATLAB script to make the displacement charts featured in this manuscript.

This work was funded by the National Institute of Health Grants (NIH/NIAID) AI103666 and Hirschl Caulier Award to GL and AI128735 to DF and GL. JM was supported by NIH MSTP training grant T32 GM7288-46 and F31 HL147470.

## Authors Contributions

JM with DF and GL input, designed, performed and interpreted most experiments, and drafted manuscript and figures. MP developed mixed BM models and cell-specific conditional knockout mice, and conducted related experiments with data interpretation. JZ helped with IVM analyses. JM, DF, GL wrote and edited the paper.

## Author Information

The authors declare that no competing interests exist.

## Supplementary Figure legends

**Figure S1, related to Figure 1.**
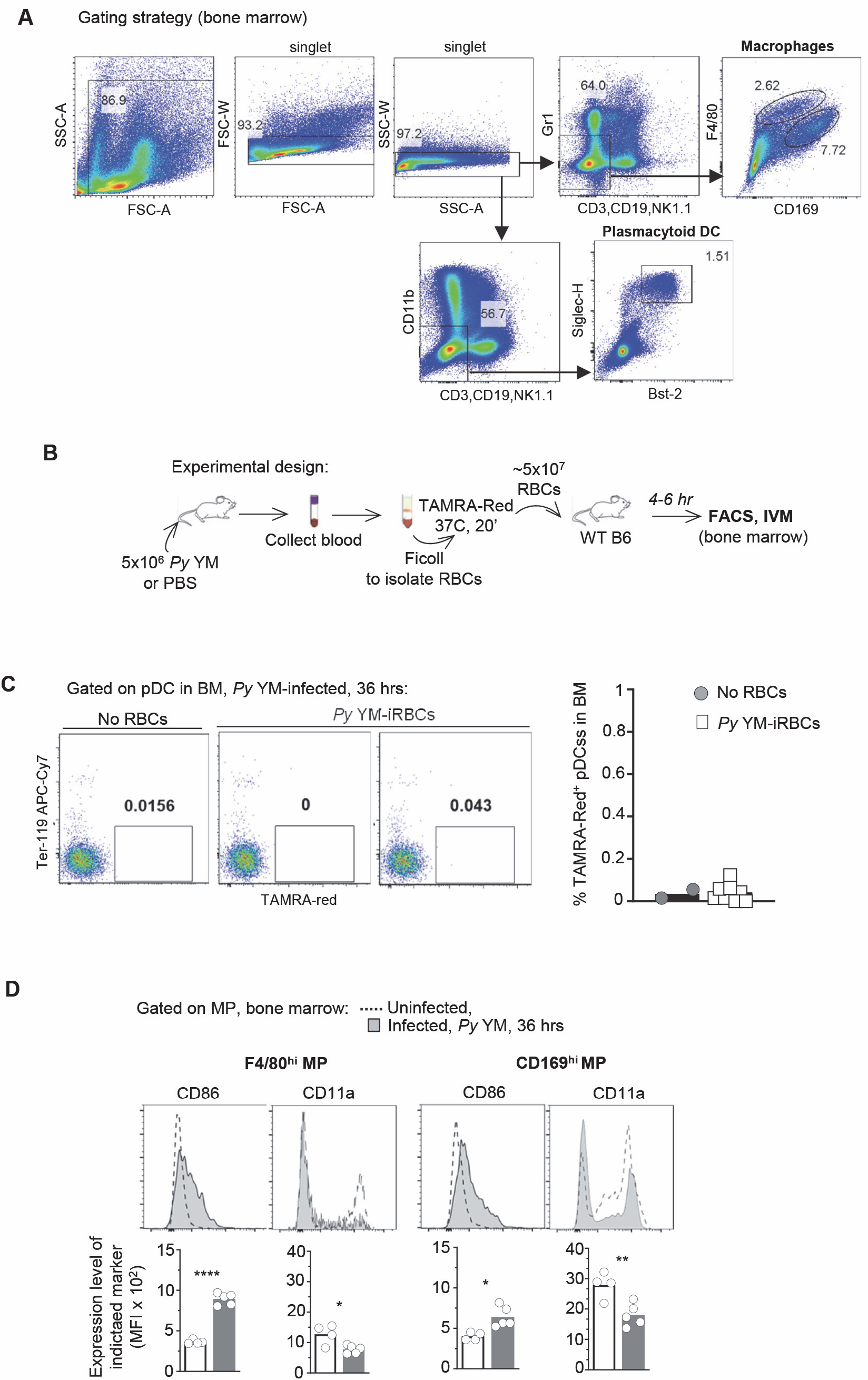
(A) Gating strategy for BM macrophages and pDC. (B) Schematic of experimental design. Peripheral blood was harvested from uninfected (uRBC) or *Py* YM-infected mice (iRBC) and labeled with the Tetramethylrodamine (TAMRA-Red) dye before transfer to naïve recipient mice. BM was harvested 6 hrs later and stained for live/dead and cell-surface expression of lin^neg^ (CD3, CD19, NK1.1), F4/80, CD169, CD11b, BST2 and SiglecH markers. (C) Analysis of TAMRA-red^+^ *Py*-iRBC pDC by FACS from experiment (B) (n=2-10 mice). (D) BM cells from uninfected or *Py* YM-infected WT mice were harvested at 36 hrs and stained with cell-surface expression of lin^neg^ (CD3, CD19, NK1.1, CD19), F4/80, CD169 and CD86 and CD11a activation markers. Representative FACS histogram overlays of CD86 and CD11a activation marker expression on CD169^hi^ and F4/80^hi^ MP with MFI across 2 pooled replicate experiments. *p*-values were calculated with \**p* < 0.05; \*\**p*< 0.01; \*\*\**p*< 0.001; \*\*\*\**p*< 0.0001; ns, not significant, using two-tailed unpaired Student’s *t* test.

**Figure S2, related to Figure 2.**
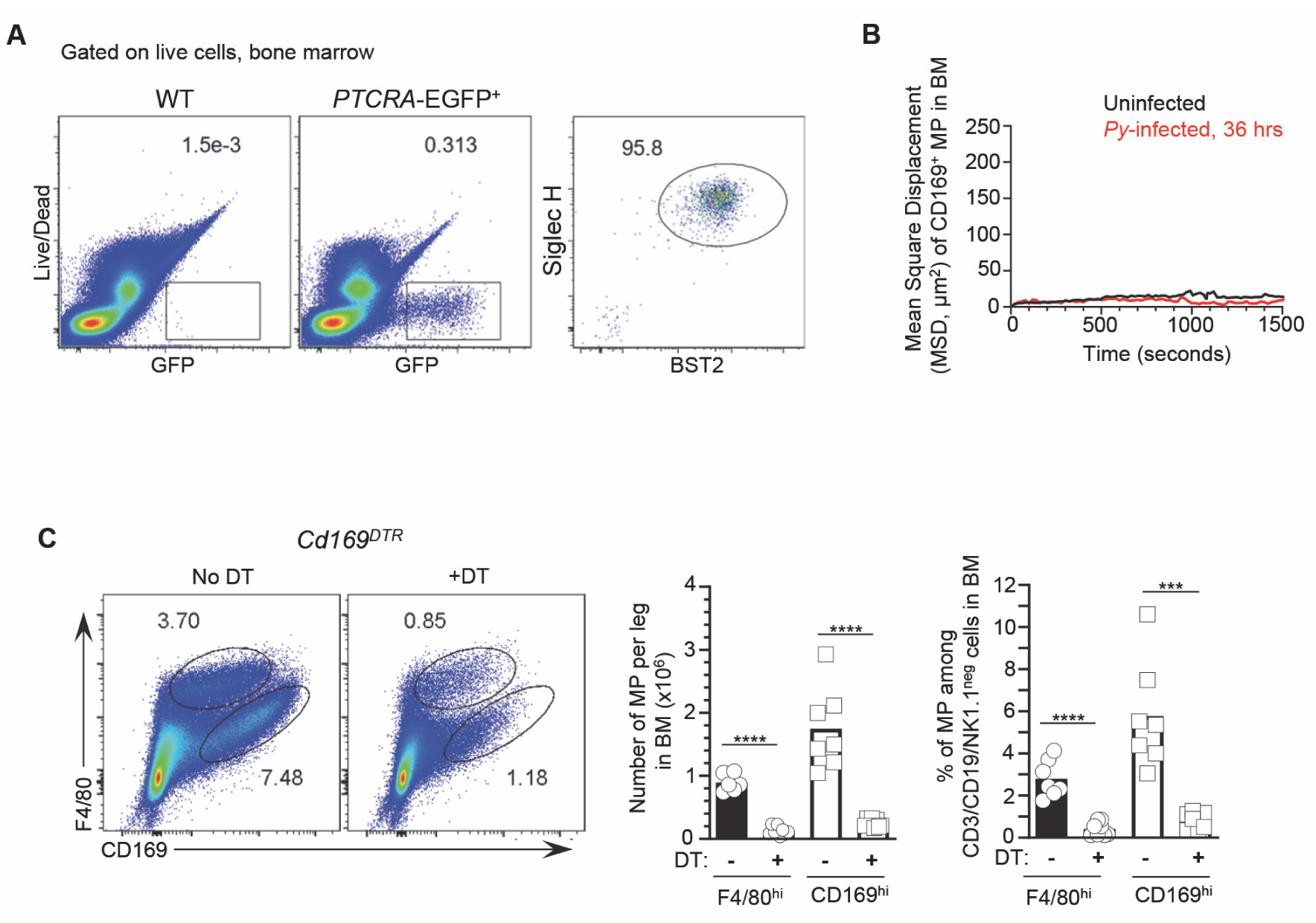
(A) BM cells from WT B6 or WT *PTCRA-*EGFP reporter mice were stained for cell-surface expression of lin^neg^ (CD3, CD19, NK1.1, CD11b, Gr1), BST-2, Siglec-H and pDC expression of GFP is shown in 1 representative of 10-15 mice. (B) MSD of CD169^+^ MP obtained after IVM imaging in the BM of uninfected and *Py*-infected WT *PTCRA-*EGFP reporter mice. (C) Representative FACS dot plots of CD169^hi^ and F4/80^hi^ MP analysis in the BM of *Cd169^DTR^*^/WT^ mice injected with DT-treated or not 12 hrs before, using cell staining as in Figure S1. Bar graphs show mean numbers of pDC per BM leg and proportion of each MP subset in 2 independent replicate experiments with each symbol representing 1 mouse (n=6 mice). *p*-values were calculated with \**p* < 0.05; \*\**p*< 0.01; \*\*\**p*< 0.001; \*\*\*\**p*< 0.0001; ns, not significant, using two-tailed unpaired Student’s *t* test.

**Figure S3, related to Figure 3.**
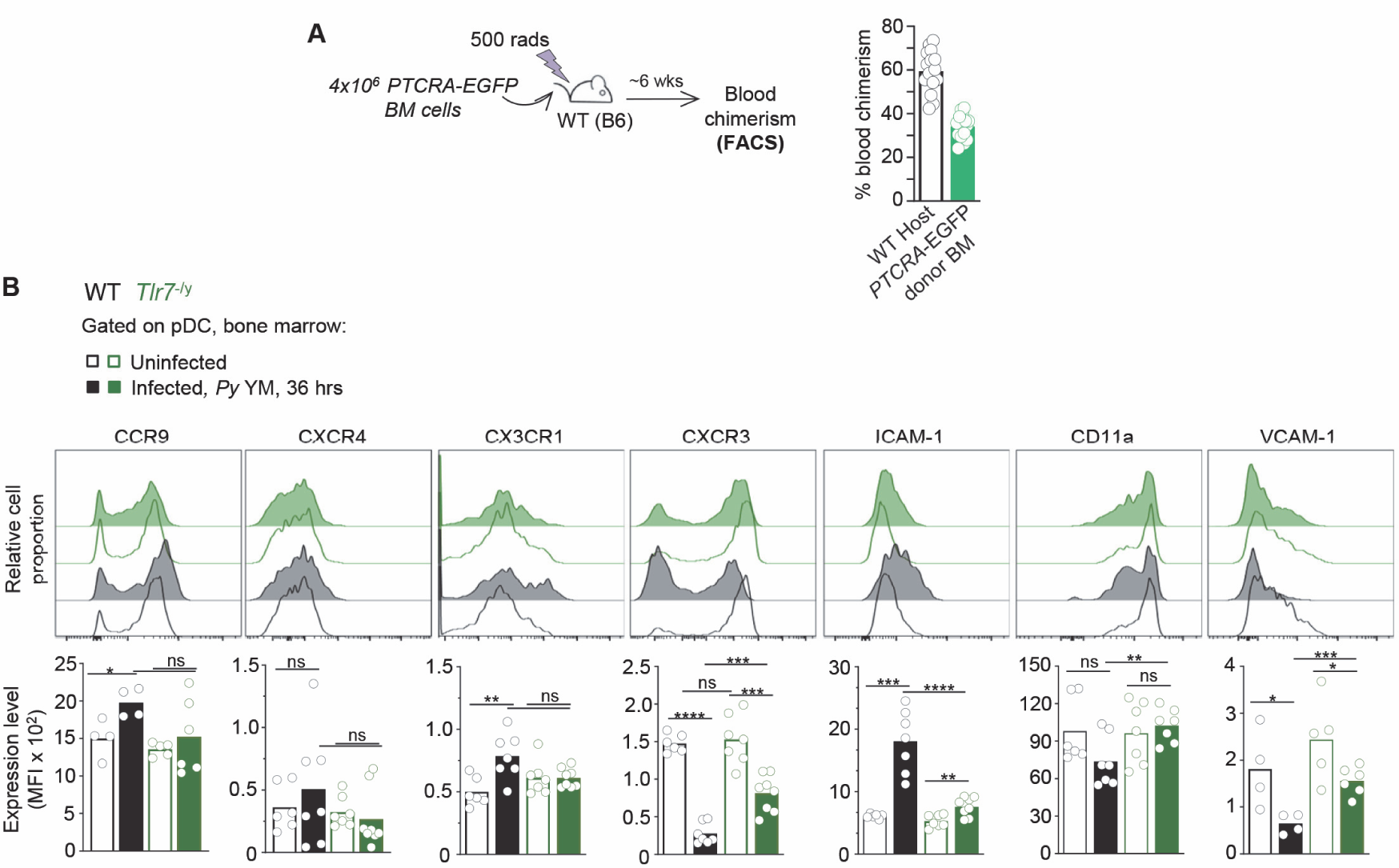
Impaired plasmacytoid dendritic cell expression of chemokine receptors and adhesion molecules in *Py*-infected TLR7-deficient mice. (A) Schematic of partial *PTCRA*-EGFP BM chimera generation and blood analysis of chimerism 6 weeks post reconstitution. (B) WT or *Tlr7^-/y^* mice either received PBS (uninfected) or were infected with 5x10^5^ *Py*-iRBCs for 36 hr before harvesting BM for FACS analysis. BM cells were stained for cell-surface expression of lin^neg^ (CD3, CD19, NK1.1, CD11b), BST-2 and Siglec-H and indicated chemokine receptors or adhesion molecules. Representative FACS histograms after gating on pDC for each experimental condition and genotype are shown. Bar graphs average cell-surface expression levels (MFI) for indicated markers across 3 replicate experiments with mean, and each symbol feature 1 mouse (n=4-8 mice)*. p*-values were calculated with \**p* < 0.05; \*\**p*< 0.01; \*\*\**p*< 0.001; \*\*\*\**p*< 0.0001; ns, not significant, using two-tailed unpaired Student’s *t* test.

**Figure S4, related to Figure 5.**
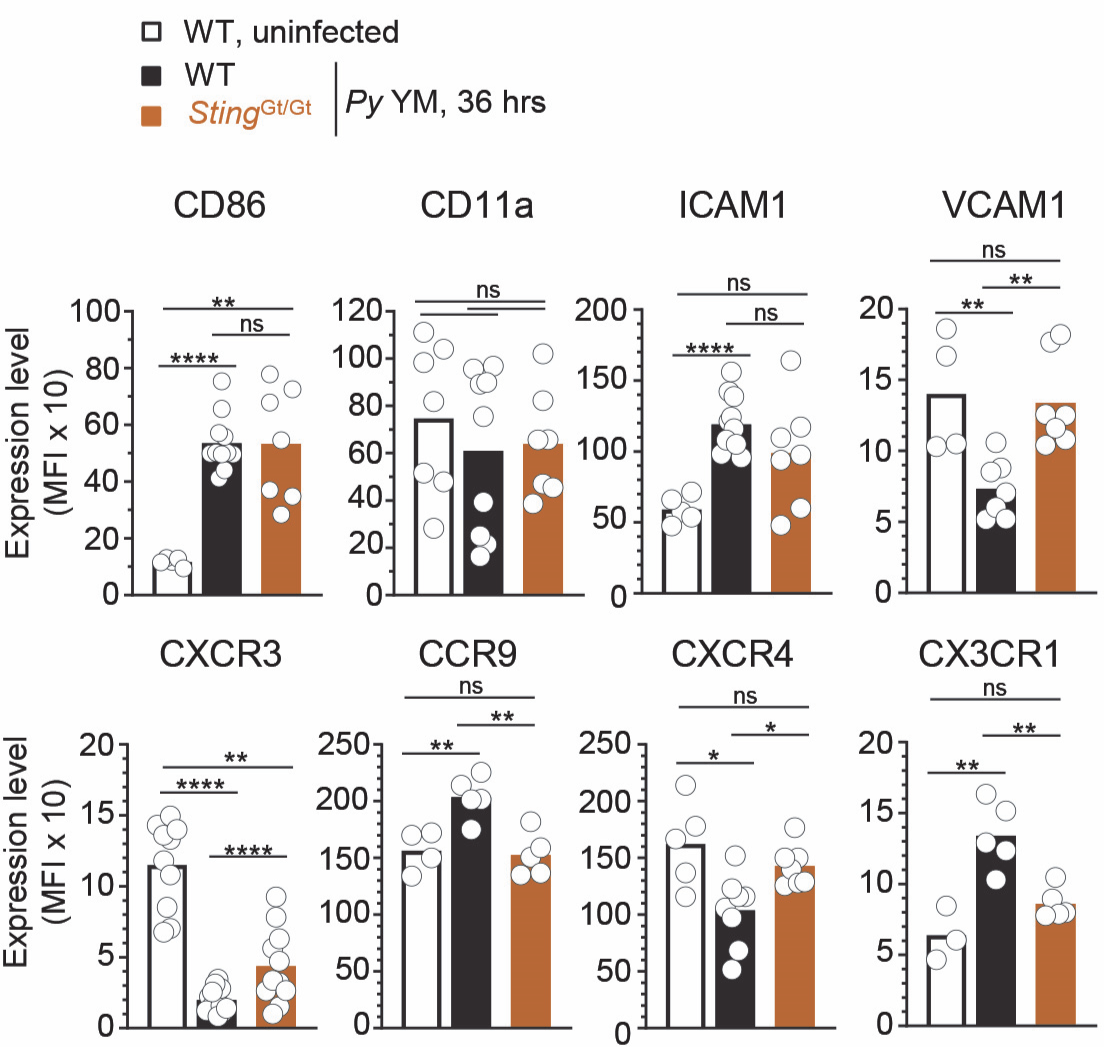
BM cells from WT B6 mice were stained cell-surface expression of lin^neg^ (CD3, CD19, NK1.1, CD11b, Gr1), BST-2 and Siglec-H and indicated chemokine receptors or no stain (Fluorescence minus one, FMO). Representative FACS histograms of 1 of 5-8 mice after gating on pDC are shown. WT- and *Sting*^Gt/Gt^-*Ifnb^YFP/YFP^* reporter mice were either injected with PBS (uninfected) or infected with 5x10^5^ *Py*-iRBCs. 36 hr later BM cells were stained with live/dead and for cell-surface expression of lin^neg^ (CD3, CD19, NK1.1, CD11b, Gr1), BST-2, Siglec-H, and indicated cell surface markers. Bar graphs quantify mean expression levels of indicated markers on pDC across 2-6 independent replicate experiments, each symbol is 1 mouse (n=3-13 mice). *p*-values were calculated with \**p* < 0.05; \*\**p*< 0.01; \*\*\**p*< 0.001; \*\*\*\**p*< 0.0001; ns, not significant, using two-tailed unpaired Student’s *t* test.

**Figure S5, related to Figure 6.**
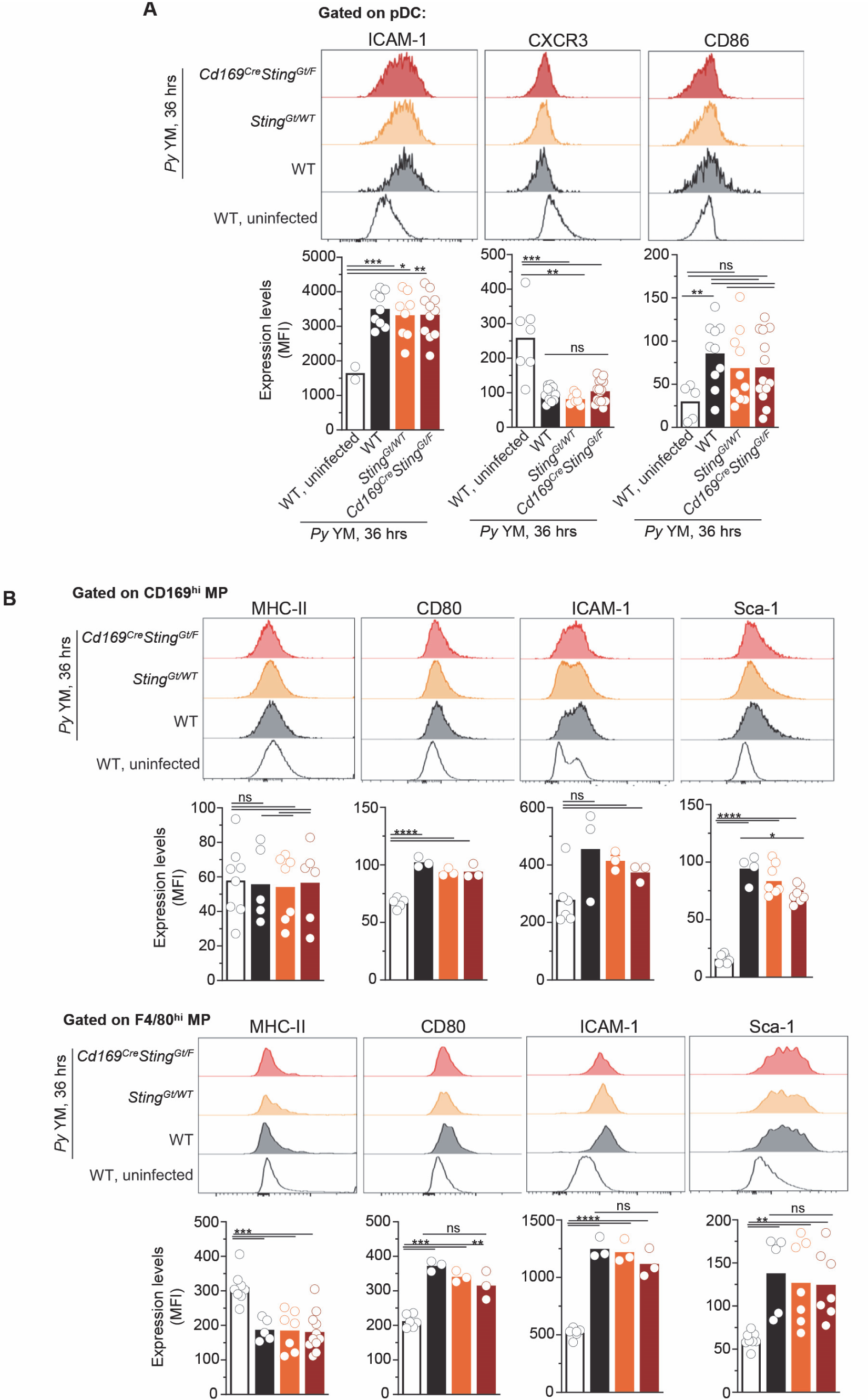
WT-, *Sting*^Gt/WT^- or *CD169^Cre/Cre^Sting^Gt/F^*-*Ifnb^YFP/YFP^* reporter mice were infected with 5x10^5^ *Py*-iRBCs. 36 hr later BM cells were stained with live/dead and for cell-surface expression of lin^neg^ (CD3, CD19, NK1.1, Gr1), CD11b, BST-2, Siglec-H, CD169, F4/80 and indicated cell surface markers. Representative FACS histograms of indicated activation markers on (A) pDC and (B) MP subsets are shown. Bar graphs pool 3 independent replicate experiments with each symbol featuring 1 mouse (n=2-18 mice). *p*-values were calculated with \**p* < 0.05; \*\**p*< 0.01; \*\*\**p*< 0.001; \*\*\*\**p*< 0.0001; ns, not significant, using two-tailed unpaired Student’s *t* test.

## Supplemental Movie legends

**Supplemental Movie 1, related to figure 1. Visualization of *Py* iRBCs uptake by CD169^+^ macrophages in the BM of living mice.** Series of movies represent 1 z plane from tibial BM of WT mice 3-5 hr after transfer of TAMRA-Red iRBC. CD169^+^ MP were visualized by injecting CD169-FITC i.v. 16 hrs prior to imaging. Movies depict iRBC flowing through vasculature, arrested in proximity with CD169^+^ MP, and moving in the BM parenchyma. Time shown in min:s.

**Supplemental Movie 2, related to figure 2. *Py* YM infection induces pDC arrest.** Visualizing pDC motility in a *PTCRA-*EGFP mouse. CD169-PE was i.v. injected 16 hrs prior to imaging to visualize CD169^+^ MP (red). pDCs (green) are highly motile in the tibial BM at steady-state (uninfected WT, left). Note that pDCs are immobile at 36 hrs post-infection with *Py* YM, exhibiting an elongated morphology in close approximation to CD169^+^ MP (red) (36h *Py* YM WT, right). Time is shown as h:mm:ss:mms.

**Supplemental Movie 3, related to figure 2. CD169^+^ MP depletion prevents pDC arrest during *Py* YM infection.** *PTCRA-*EGFP*/Cd169^DTR/WT^* mice were treated with control PBS (left) or diphtheria toxin (DT) (right) 12 hrs prior to *Py*-infection. 36 hrs post-*Py* YM infection pDC (green) motility is significantly increased in DT treated mice (right) versus vehicle treated controls (left). Time is shown as h:mm:ss:mms.

**Supplemental Movie 4, related to figure 3. pDCs are highly motile in BM of WT and *Tlr7*^-/y^ naïve mice.** Four-dimensional data series represented using extended focus from tibial BM of WT or *Tlr7^-/y^ PTCRA-*EGFP mice at steady-state. CD169^+^ MP were visualized by injecting CD169-PE i.v. 16 hrs prior to imaging. EGFP-pDCs are shown in green. Time is shown as h:mm:ss:mms. Scale bar represents 20 μm.

**Supplemental Movie 5, related to figure 3. pDC remain highly motile in the BM of *Tlr7*^-/y^ mice during *Py* YM infection.** Four-dimensional data series represented using extended focus from tibial BM of WT or *Tlr7^-/y^ PTCRA-*EGFP N1 mice that had been infected with *Py* YM 36 hrs prior to imaging. CD169^+^ MP were visualized by injecting CD169-PE i.v. 16 hrs prior to imaging. EGFP-pDCs are shown in green. Time is shown as h:mm:ss:mms.. Scale bar represents 40 μm.

**Supplemental Movie 6, related to figure 3.** TLR7-deficient pDC fail to arrest in the BM of *Tlr7^-/y^ PTCRA-*EGFP/WT partial BM chimeras during *Py* YM infection.

Intravital imaging in the tibia demonstrates pDC (green) arrest in *PTCRA-*EGFP/WT (left) but not *Tlr7^-/y^ PTCRA-*EGFP/WT (right) chimeras 36h post-infection with *Py* YM. To visualize CD169^+^ MP, mice were injected with CD169-PE i.v. 16 hrs prior to imaging. Scale bar represents 50 μm. Time is shown as h:mm:ss:mms.

**Supplemental Movie 7, related to figure 4. Blocking G_α_I-signaling induces robust pDC arrest and a morphologically distinct phenotype.** WT *PTCRA-*EGFP reporter mice either received PBS or pertussis toxin (PT) and were infected with 5x10^5^ *Py*-iRBCs. To visualize CD169^+^ MP, CD169-PE mAb was i.v. administered 16 hrs prior to IVM of tibial BM. Intravital imaging in the tibia demonstrates pDCs (green) arrest 36 hrs post-*Py* YM in vehicle and PT treated mice. Note that pDC morphology is distinctly rounded in setting of PT treatment (right), although directly adjacent to MP (red) in both preps. Scale bar represents 30 μm. Time is shown as h:mm:ss:mms.

**Supplemental Movie 8, related to figure 5. pDC movement and clustering in the BM of naïve compared to *Py* YM-infected *Sting^Gt/Gt^* mice.** Four-dimensional data series represented using extended focus from tibial BM of *Sting^Gt/Gt^ PTCRA-*EGFP N9 mice at steady-state or 36 hrs post-infection *Py* YM. CD169^+^ MP were visualized by injecting CD169-PE i.v. 16 hrs prior to imaging. EGFP-pDCs are shown in green. Time is shown as h:mm:ss:mms. Scale bar represents 40 μm.

**Supplemental Movie 9, related to figure 5. WT pDCs arrest and form large clusters in *Py* YM-infected *Sting^Gt/Gt^ PTCRA-*EGFP/WT chimeras.** WT *PTCRA-*EGFP/WT partial BM chimeras were infected with *Py* YM. 36 hrs post-infection, intravital microscopy imaging of the BM reveals markedly increased quantity and side of pDC (clusters) in *Sting^Gt/Gt^ PTCRA-*EGFP/WT (left) compared to WT *PTCRA-*EGFP/WT (right) chimeras. Time is shown as h:mm:ss:mms. Scale bar represents 50 μm.

**Supplemental Movie 10, related to figure 5. pDC form large clusters in *Py* YM-infected *Sting^Gt/Gt^* compared to WT mice.** Four-dimensional data series represented using extended focus from tibial BM of WT or *Sting^Gt/Gt^ PTCRA-*EGFP N9 mice 36 hrs post-infection *Py* YM. CD169^+^ MP were visualized by injecting CD169-PE i.v. 16 hrs prior to imaging. EGFP-pDCs are shown in green. Clusters of pDCs around CD169^+^ MP are emphasized with a dotted line encircling them. Yellow arrowheads pointed to the relatively sparse pDCs in WT compared to *Sting^Gt/Gt^* mice. The lucent tissue in RL movie is bone. Time is shown as h:mm:ss:mms. Scale bar represents 40 μm.

